# Sustained Strain Applied at High Rates Drives Dynamic Tensioning in Epithelial Cells

**DOI:** 10.1101/2024.07.31.606021

**Authors:** Bahareh Tajvidi Safa, Jordan Rosenbohm, Amir Monemian Esfahani, Grayson Minnick, Amir Ostadi Moghaddam, Nickolay V. Lavrik, Changjin Huang, Guillaume Charras, Alexandre Kabla, Ruiguo Yang

## Abstract

Epithelial cells experience long lasting loads of different magnitudes and rates. How they adapt to these loads strongly impacts tissue health. Yet, much remains unknown about the evolution of cellular stress in response to sustained strain. Here, by subjecting cell pairs to sustained strain, we report a bimodal stress response, where in addition to the typically observed stress relaxation, a subset of cells exhibits a dynamic tensioning process with significant elevation in stress within 100s, resembling active pulling-back in muscle fibers. Strikingly, the fraction of cells exhibiting tensioning increases with increasing strain rate. The tensioning response is accompanied by actin remodeling, and perturbation to actin abrogates it, supporting cell contractility’s role in the response. Collectively, our data show that epithelial cells adjust their tensional states over short timescales in a strain-rate dependent manner to adapt to sustained strains, demonstrating that the active pulling-back behavior could be a common protective mechanism against environmental stress.

## INTRODUCTION

Epithelial cells are subjected to strains of varied rates and magnitudes^1, 2, 3^. Some deformations are sustained for a period of several minutes rather than acute lasting only seconds. Cells have effective mechanisms, such as stress relaxation, to dissipate stress and prevent tissue damage^4, 5^. Stress relaxation is often achieved through active remodeling of biopolymers in cells, which has been previously shown in a monolayer study by Khalilgharibi et al^5^. This remodeling helps to alleviate internal stress but also allows the tissue to accumulate large cellular deformations by creep if the tension is sustained over time. Strain stiffening is one of the effective mechanisms tissues employ to limit the strain cells are subjected to. Active contraction also provides a way to limit deformations; it occurs in the muscle where individual myofibers exhibit stiffening behavior by adjusting myosin attachment kinetics under load^6^. However, the generality of such behavior is unclear, and it is unknown if cells in epithelia can respond actively when subjected to high strain-rate, sustained deformation. During physiological function, epithelial cells experience strains applied at widespread rates. Therefore, the timescale of the strain application may be an important factor in determining the response outcome^7^. For instance, force application rate determines the activation of talin-dependent mechanosensing^7^. We have shown that cellular response to ramp strains is rate dependent, with increasing stress levels and higher probability of cell pair fracture at higher strain rates^8^. However, the role of strain rate in determining cellular response to sustained strain is still not fully understood.

Cells and tissues display two broad categories of responses to deformation: a fast response, that involves primarily the cytoskeletal structures already present within the cell, and a slower response, that necessitates reorganization of the cytoskeleton downstream of signaling, transcription, cell division, etc. The first response can be essentially viewed as a safety mechanism that counters transient mechanical overload, while the second aims to adapt to chronic overload. The second only comes into play in response to repeated or sustained overload.

Currently, our understanding of how cells and tissues adapt to this sustained strain is limited. It is unclear whether the cellular response is cell autonomous or stems from the interactions between neighboring cells in a tissue.

Most of the previous efforts to explore the adaptation mechanisms focus primarily on cytoskeleton reorganization resulting from the mechanical loading, which is evaluated using imaging techniques^9, 10, 11, 12, 13^. Some studies also delve into the mechanical responses of these systems, including stress and strain, although these efforts are largely limited to cell monolayers or single cells. For instance, previous experiments examining the response of suspended monolayers to deformation report stress relaxation mediated by turnover of the actin cytoskeleton but no active response^5, 14^. Yet, other studies point to active responses of cells when they are adherent and subjected to force^15^. Thus, focal adhesions may represent a key element in sensing mechanical stress.

Since cell-cell junctions are one of the primary components of tissues that enable them to withstand mechanical loading, any perturbations in these intercellular molecules, caused by mutations or diseases, could jeopardize tissue integrity and potentially lead to organ failure. Consequently, understanding the influence of physiologically relevant mechanical loading on biophysical characteristics of cellularized materials at the cell pair scale has a substantial clinical relevance. Yet it is underexplored largely due to the lack of a proper method to allow precise displacement application and force measurement when cells are in their physiological state. Here, we will use our previously developed single cell adhesion micro tensile tester (SCAμTT) device^8^ to examine the respective contributions of intercellular adhesions and focal adhesions to the safety mechanism that enables cells to withstand transient mechanical load. To this end, we subject adherent epithelial cell pairs or single cells to a defined strain and monitor their stress response. This allows us to track stress within the individual cells and investigate the mechanism of adaptation during stress response. Strikingly, we find that epithelial cells indeed exhibit active tensioning in a strain and strain-rate dependent manner in response to a sustained strain. Stress relaxation dominates cellular response at low strain rates during holding, while high strain rates increase the probability of triggering the dynamic tensioning response. This tensioning response depends on actomyosin and is observed in both single cells and cell doublets, suggesting it may be sensed by focal adhesions.

## RESULTS

### Cells increase their tension in response to a high strain rate deformation

To investigate the change in stress in a cell pair in response to the application of strain, we used our SCAμTT platform (Fig. 1a)^8^, which comprises two islands separated by a negligible gap following the printing process while keeping the two islands mechanically separated. Cell pairs seeded onto the platform adhere via focal adhesions and form intercellular junctions that span the gap between the two islands. The device allows stretching a cell pair while measuring applied stress and strain. Briefly, a displacement of D imposed on the forcing beam (Island 2) induces a deflection of δ in the sensing beam (Island 1) via forces transmitted across the cell-cell contact (Fig. 1b). Based on beam deflection, force and displacement in the cell pair can be calculated (see Methods). Importantly, the stiffness of the beams is set such that δ is much smaller than D, and therefore the deformation of the cell pair is assumed to be determined by D alone^16^. Therefore, by controlling the rate at which D is applied, we can control the rate at which the cell pair is deformed (Fig. 1c).

**Figure 1.**
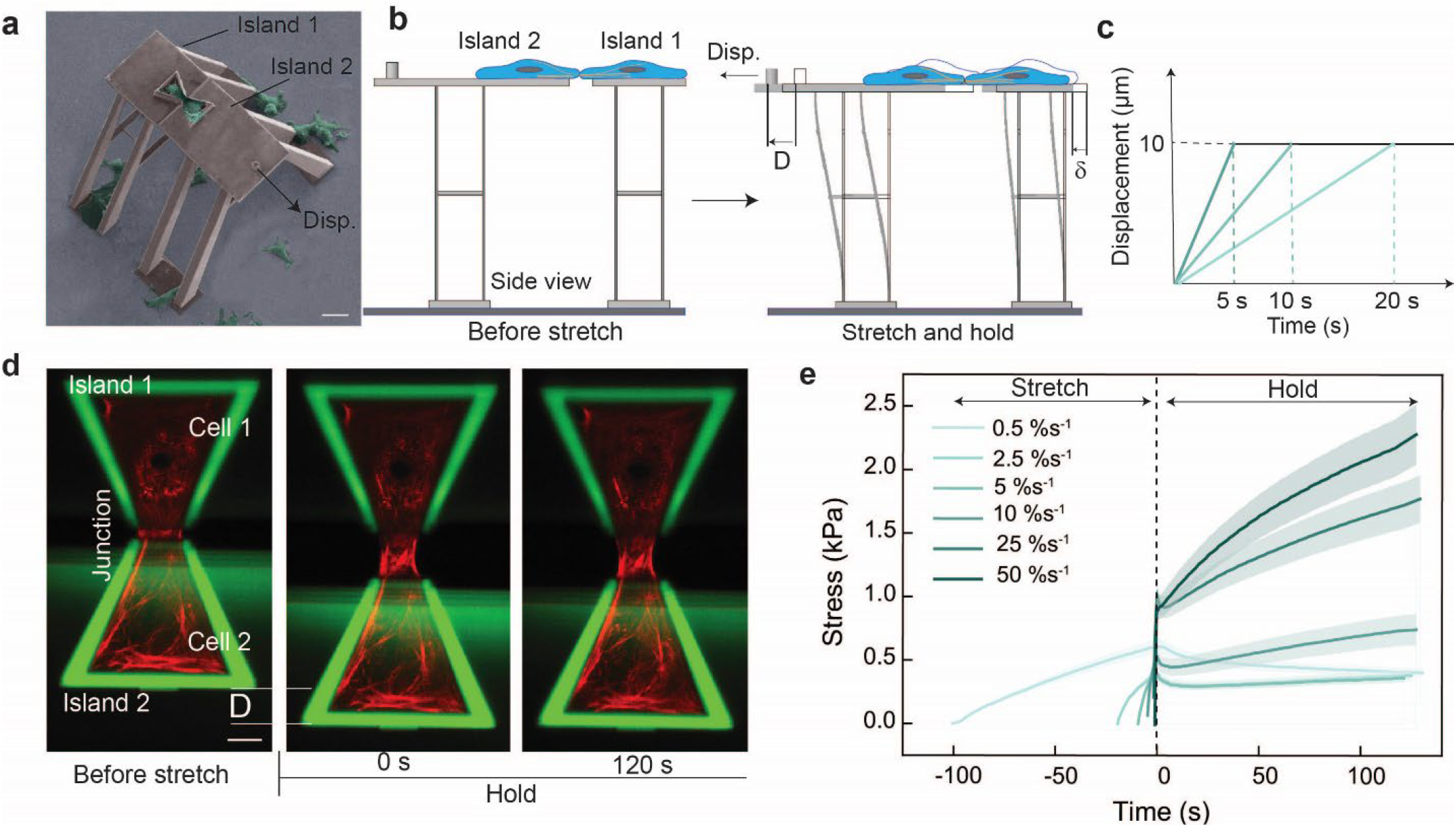
Mechanical response of epithelial cell pairs is strain rate dependent in ramp and hold experiments. **a.** SEM image of the single cell adhesion micro tensile tester platform used to stretch cell pairs. Scale bar: 50 μm. **b**. Schematic diagram presenting the method used to apply strain to the cell pair. Cells (blue) were deposited on each part of the bowtie and had time to make mature junctions. Then, they were stretched at different rates, and using the deflection in island 1 (δ) and island 2 (D), the stress and strain in the cell pair was calculated. **c**. Time course of the applied 50% strain with different rates. **d**. Confocal images of the cells prior and post stretch application (red: F-actin; green: structure autofluorescence). The cells are stretched at 25 %s^−1^. Scale bar: 10 μm. **e**. Temporal evolution of stress (average of all curves for each condition ± s.e.) for different rates. To make the comparison easier to visualize, we aligned the initiation of the holding phase to t = 0 s.

The details of stress and strain definition are provided in the Methods section and our previous work^8^. Briefly, engineering stress is calculated by dividing force by the initial cross-sectional area of the cells, which is determined by multiplying the width and height of the cell-cell junction. Both dimensions are measured using fluorescence images^8^. Actin and intermediate filaments serve as the main structures that transmit force between cells via intercellular adhesions and their number is expected to remain constant in our experiments^8^. Although the junction’s area may vary slightly during loading, the linker density would change accordingly. This is why we decided against normalizing the force by the current area and instead used a constant cross-sectional area in our stress calculations. Consistently with this decision, we defined strain as displacement over the initial length of the cell pair. To identify the relevant material length, we carefully examined our recorded videos and observed that the regions near the junction deformed the most, while the areas beyond the nucleus remained unaffected by the external load^*8*^. Therefore, we defined the initial length of the cell pair as the nucleus-to-nucleus distance.

In this study we investigated the response of A431 epithelial cells to a ramp and hold condition, in which a predetermined strain is applied at a controlled rate and then held constant. Under load, tension is applied to both cell-cell and cell-ECM adhesions as the cytoskeleton is deformed, actively responds, and adapts to the loading environment (Fig. 1d). We first subjected cell pairs to a strain of 50% at strain rates varying from 0.5 %s^−1^ to 50 %s^−1^. These conditions are within the physiological range (0.5-80 %s^−1^)^7^ and similar to those previously used in monolayer studies^5, 17, 18^. Surprisingly, the temporal evolution of stress is quantitatively dependent on the applied strain rate (Fig. 1e, Supplementary Movie 1).

Depending on the strain rate, characteristics of these stress versus time curves are different. Stress first increased with the ramp deformation and peaked when the target deformation was reached (defining t = 0 s). Consistent with the known viscoelastic behavior of cells, peak stress increased from just below 0.5 kPa to 1 kPa with increasing strain rate. Then, regardless of the strain rate, stress briefly relaxed by about 100 Pa following a power law in the first ~10s (Supplementary Fig. 3c-d), possibly due to passive viscoelastic behavior and remodeling of the cytoskeleton, as previously described by others^19^. After this short dip in stress, the temporal evolution of stress in the following 2 minutes depended greatly on the applied strain rate, displaying a bimodal behavior. For strain rates ≤5 %s^−1^, the stress remained approximately constant at the stress achieved after the relaxation phase, indicative of a solid-like behavior. However, for strain rates ≥10 %s^−1^, in some cell pairs, the stress rose linearly (i.e., tensioning) reaching values larger than the stress at t = 0 s (Fig. 1e, Supplementary Fig. 2, 2d-f). After this linear phase in the first 30 to 50 seconds, stress started to trend towards a new steady state (Fig. 1e). At the end of the two-minute observation, in cells which were deformed at 10 %s^−1^, the stress increased by ~40% from 0.5 kPa at t = 0 s to 0.7 kPa. For 25 %s^−1^ and 50 %s^−1^, the stress approximately doubled. Overall, the rate of stress enhancement increased with strain rate. In conjunction with the stress increase, we also observed a decrease in strain at t = 2 min (as shown in Supplementary Fig. 3b).

These results suggest that cells are actively reacting to the loading conditions and exhibit varied responses dependent on those conditions. When the strain rate is low, the cells relax stress but when the strain rate is high, they appear to exert more tension, perhaps aiming to reduce the deformation that they are subjected to.

### Characterizing stress accumulation in cell pairs

Pooling individual curves for each condition revealed two distinct behaviors in cell pairs subjected to high strain rates (10 %s^−1^, 25 %s^−1^ and 50 %s^−1^). In some cell pairs, the stress relaxes and reaches a plateau after a few seconds, whereas, for others, the relaxation phase is followed by a linear increase in stress, i.e., tensioning (Fig. 2a, Supplementary Fig. 2, Supplementary Movie 2).

**Figure 2.**
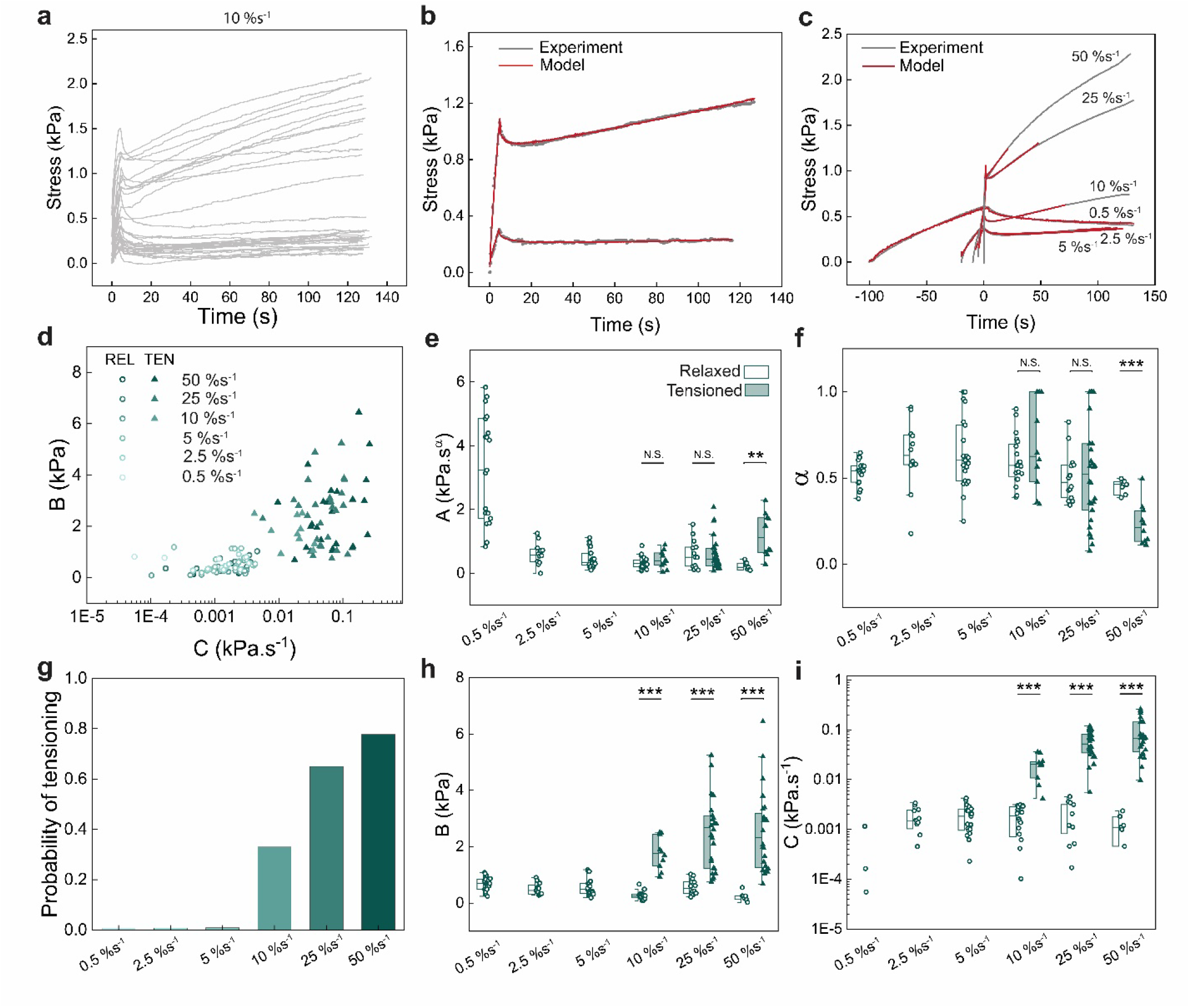
At 50% strain, the mechanical response of the cell pair to high strain-rate stretch is bimodal: relaxed and tensioned. **a.** Temporal evolution of the stress in the cell pair following a 50% strain applied at 10 %s^−1^ (N = 30). **b**. The example experimental relaxed and tensioned curves (gray) are fitted with the empirical model (red). **c**. Average stress-time curves obtained in ramp-hold experiments for various levels of strain rate (gray) are fitted with the empirical model (red). To make the comparison easier to visualize, we aligned the initiation of the holding phase to t = 0 s. **d**. B vs C values for all strain rates applied to reach 50% strain are plotted and clustered on a semi logarithmic scale. The results of clustering are represented by solid triangles and hollow circles to represent tensioned and relaxed curves, respectively. **e-f, h-i**. Box plots representing parameters *A* · *ε*_0_ (amplitude of relaxation), *α* (power law exponent), *B* · *ε*_0_ (residual stress), and *C* · *ε*_0_ (captures the slope of tensioning) for the cell pairs experiencing various levels of strain rate. **g**. Probability of tensioning for each strain rate, which was calculated by dividing the number of tensioned curves by the number of all curves for each condition.

To capture the mechanical characteristics of the stress response, we fitted our experimental stress curves with an empirical fit function for the relaxation modulus of the material, *G*(*t*) = *At*^−*α*^ + *B* + *Ct*, based on our characterization of the response to a step strain *ε*_0_ (Fig. 2b, 2c and Supplementary Fig. 3, Methods). We do not assign any physical meaning to the parameters extracted from fitting the experimental data with G(t) and only use this function to mathematically describe the experimental data. This approach assumes that the response is linear with the deformation for the purpose of quantifying the material’s response to arbitrary loads. The power law function defines the first relaxation phase, where *A* · *ε*_0_ sets the amplitude of stress relaxation and *α* is the power law exponent, *B* · *ε*_0_ is the residual stress in the relaxed curves and an estimation of the stress in the transition point between the relaxed and tensioned part of the response for tensioned curves, and *C* · *ε*_0_ captures the slope of the tensioning part of the response (a linear fit for C is shown in Supplementary Fig. 3f). To categorize the two visually distinguishable groups in the stress response, we used the values of B and C which serve as indicators of the second phase of the response and exhibit marked differences between the relaxed and tensioned curves. First, we used an unbiased approach by plotting these two parameters on a semilogarithmic scale and then clustering our experimental data based on the characteristics of the second phase of the stress response (Methods). Remarkably, all the cell pairs subjected to low strain rates (0.5 %s^−1^, 2.5 %s^−1^ and 5 %s^−1^) exhibit a relaxed response (Fig. 3d, hollow circles), whereas the cell pairs that experienced high strain rates (10 %s^−1^, 25 %s^−1^ and 50 %s^−1^) can be categorized into two groups, i.e., relaxed and tensioned, consistent with our qualitative interpretation (Fig. 3d, and Supplementary Fig. 1) and independent of the optimization and clustering algorithm used (Supplementary Fig. 4).

**Figure 3.**
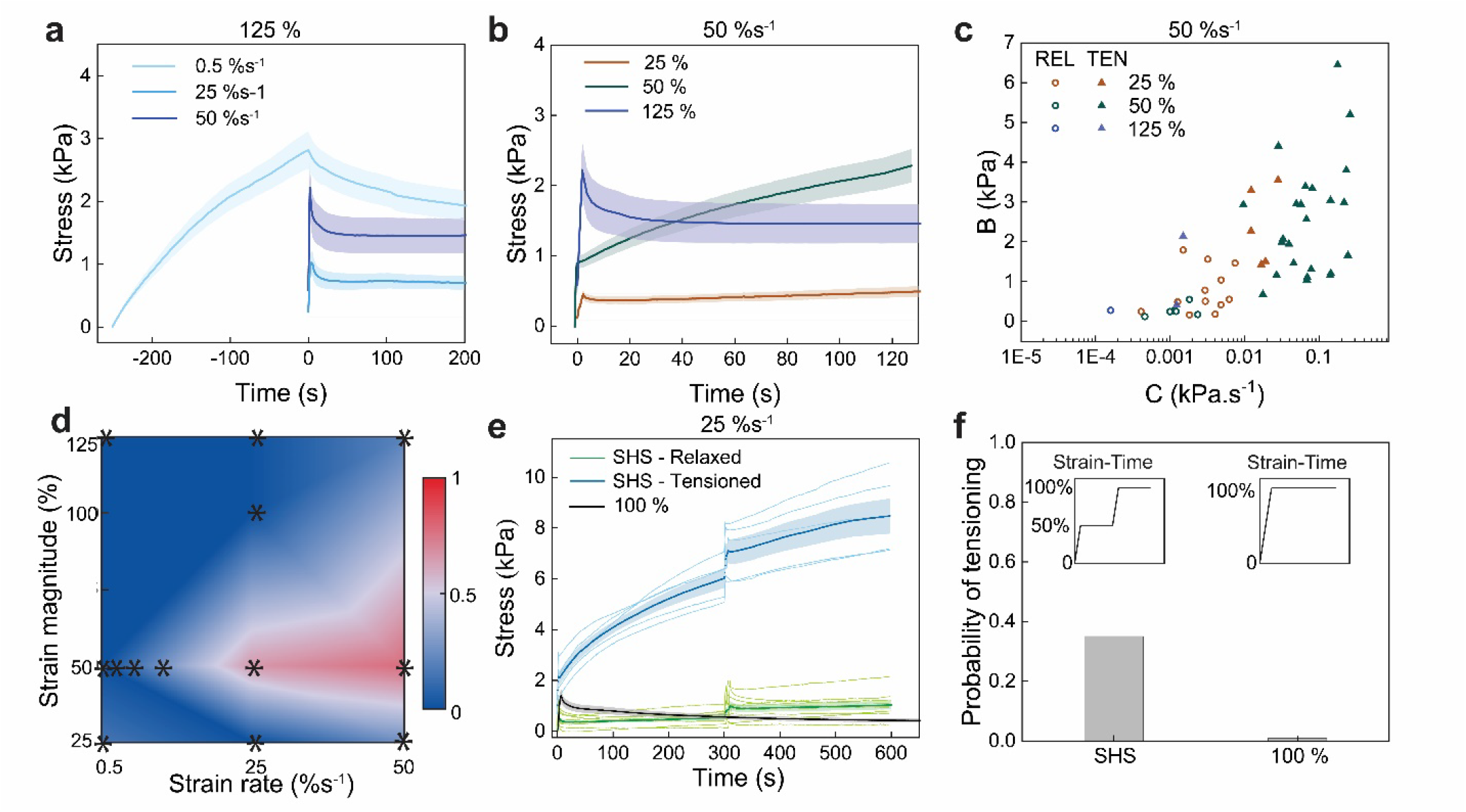
The effect of the strain magnitude on the mechanical response of cell pairs subjected to high strain rates. **a.** Temporal evolution of stress (average of all curves for each condition ± s.e.) over time for cell pairs stretched at different rates to reach 125% strain. To make the comparison easier to visualize, we aligned the initiation of the holding phase to t = 0 s. **b**. Temporal evolution of stress for cell pairs stretched at a strain rate of 50 %s^−1^ to reach various strain magnitudes. **c**. B vs C values calculated from fitting the temporal evolution of stress over time in response to different strain magnitudes applied at 50 %s^−1^ are plotted and clustered on a semi logarithmic scale. Solid triangles and hollow circles represent tensioned and relaxed responses, respectively. **d**. The probability of tensioning versus the strain magnitude and strain rate. The stars indicate the probability of tensioning calculated from the experimental results. **e**. Temporal evolution of the stress in the cell pair in response to stretch-hold-stretch (N = 14), and a single stretch to 100% strain (N = 12). In the stretch-hold-stretch experiments, the cell pairs were stretched to 50% at 25 %s^−1^ strain rate. After 5 minutes of hold, the cell pairs were stretched at 25 %s^−1^ to reach a total of 100% strain. **f**. Probability of tensioning for stretch-hold-stretch experiments and a single stretch to 100% strain. The inset shows the applied strain over time for a stretch-hold-stretch experiment and a single stretch to 100% strain.

Based on our classification, the initial stress relaxation captured by the power law was similar for both categories, as the distributions of *α* and A were not different between tensioning and relaxing cell pairs, except for strain rates of 50 %s^−1^ (Fig. 2e, f, Supplementary Fig. 5). At low strain rates, cells displayed a power law exponent *α*~0.5, larger and more fluid-like than previously reported for the cortex of single cells^20^ or for epithelia devoid of a substrate^5^. This difference may arise because our experiments probe regions of the cell interfaced to the substrate. At 50 %s^−1^ for cells that displayed tensioning behavior, *α* was significantly lower, reaching a value of 0.23 ± 0.12, suggesting that some reinforcement may occur at the highest strain rates to render the cells more solid-like and prevent flow.

When we examined the second part of the response, we found that B and C were significantly larger for tensioning cells than for relaxing cells subjected to the same strain rate (Fig. 2h-i). This indicated that cells with higher B, which represents the elasticity of the system, initiate tensioning in response to high strain rates. We next investigated a potential relationship between B and C. While we could not find any correlation for cells that relaxed (Pearson correlation r = 0.05), a positive correlation was detected for cells with a tensioning response (Pearson correlation r = 0.34) (Fig. 2d). In addition, when we computed the probability of tensioning for a given strain rate, we found that this increased with strain rate (Fig. 2g and Supplementary Table S1).

### Tensioning is suppressed by large deformations

We next investigated the dependency of the tensioning behavior on the applied strain magnitude. The strain magnitude was increased to 125%, a strain level that epithelial cells such as keratinocytes can tolerate without losing integrity or viability^21^, and strain rates of 0.5 %s^−1^, 25 %s^− 1^, and 50 %s^−1^ were used. Interestingly, for this large strain, the average stress curve shows predominantly a relaxation behavior across these strain rates (Fig. 3a). We theorized that high strain applied in one step could result in cytoskeleton damage and thus prevent active tensioning responses and therefore compared the behavior of cell pairs stretched at 50 %s^−1^ subjected to strains of 25%, 50%, and 125% (Fig. 3b). Qualitatively, tensioning appears to dominate at 50% strain, but for both 25% and 125% strain, cell pairs appear to predominantly exhibit a relaxing behavior. To understand this phenomenon further, we plotted B vs C and clustered our data to determine the probability of tensioning and relaxing for each strain (Fig. 3c). This reveals that 77% of cell pairs stretched to 50% display tensioning compared to just 25% and 20% for strains of 25% and 125%, respectively, suggesting the existence of a functional strain range of the response (Supplementary Fig. 7e-g, and Supplementary Table S1). The probability of tensioning for all tested combinations of strain and strain rate are shown in Fig. 3d to visualize the combined influence of these parameters on tensioning (Supplementary Table S1, see Supplementary Fig. 6-8 for summary of all data). The plot shows that tensioning is most likely to occur at an intermediate strain magnitude of 50% at strain rates above 25 %s^−1^. Tensioning is unlikely to occur at low strain rates regardless of the applied strain, as well as at the low and high ends of the strain magnitude. We also explored a potential role of stress threshold in activating the tensioning response by producing a plot similar to that in Fig. 3d. For this, we plotted the normalized average values of stress at the end of loading for each condition and found that this plot does not resemble the probability of tensioning plot (Supplementary Fig. 9). Furthermore, no correlation was found between the two plots (Pearson correlation r= −0.09). Therefore, we concluded that the tensioning response is not triggered by stress at the end of the loading phase.

To further investigate the hypothesis that high strains applied at high rate may suppress tensioning behavior due to actin disruption, we conducted a stretch-hold-stretch experiment. In these, a total strain of 100% was applied to the cell pairs at a rate of 25 %s^−1^, but strain was subdivided into two steps of 50% with a hold time of 5 minutes between the steps (Fig. 3e). We reasoned that such a pause may provide cells with sufficient time to remodel their cytoskeleton and prevent damage because the duration of the pause is commensurate with the characteristic turnover time of actomyosin^5, 22^. Consistent with our initial experiments (Fig. 1), tensioning behavior is seen in some cells in the first step of the experiment. In the second step of the experiment, cell pairs that exhibited tensioning in the first step continued with this behavior, while those that initially relaxed continued to relax. In contrast, when the 100% strain is applied in a single step, all cell pairs exhibit relaxing behavior (Fig. 3e, f and Supplementary Table S1). These results support the idea that the tensioning behavior can be observed for larger strains when cell pairs are allowed to respond and adapt, and that application of large, sudden strains may lead to damage that prevents adaptation.

### Tensioning depends on actomyosin contractility and is not triggered by the intercellular adhesion molecules

Next, we investigated the molecular mechanisms underlying sensing of strain and transduction of the tensioning response. In epithelial cells, mechanosensitive processes have previously been reported to take place at focal adhesions^23, 24^ and at intercellular junctions^25^. To elucidate the role of intercellular junctions in sensing the applied strain, we modified our platform such that a single cell could bridge the gap between islands. Strikingly, single cells displayed tensioning responses similar to cell pairs (Supplementary Fig. 10), indicating that intercellular adhesions are not necessary to sense strain and trigger tensioning.

As actomyosin plays a key role in generating the substrate stress in single cells and cell pairs^26, 27, 28^, we investigated its role in controlling the tensioning behavior. We first examined how F-actin organization was influenced by the applied strain rate by tracking its localization and intensity with a live stain for 10 minutes after stretching the cells to 50% strain at a strain rate which does not lead to tensioning (0.5 %s^−1^) and at one that does (50 %s^−1^) (Fig. 4a). Further, as a control in which active responses are abrogated and to account for changes in fluorescence intensity due to imaging-induced photobleaching, pairs of cells fixed with paraformaldehyde were also stretched at both rates (Supplementary Movie 3). In response to stretch, F-actin intensity in live cell pairs increased in the confinement region relative to the fixed control for both strain rates, although the increase was larger at 50 %s^−1^ (Fig. 4b (i)). Near the cell-cell junction, there was no difference between the live cells stretched at either rate although the intensity did increase relative to the fixed controls (Fig. 4c (i)). These results support the previous findings that the cell-cell junction might not be required for triggering the tensioning behavior and support the idea that reorganization of the actin cytoskeleton occurs concomitantly with tensioning.

**Figure 4.**
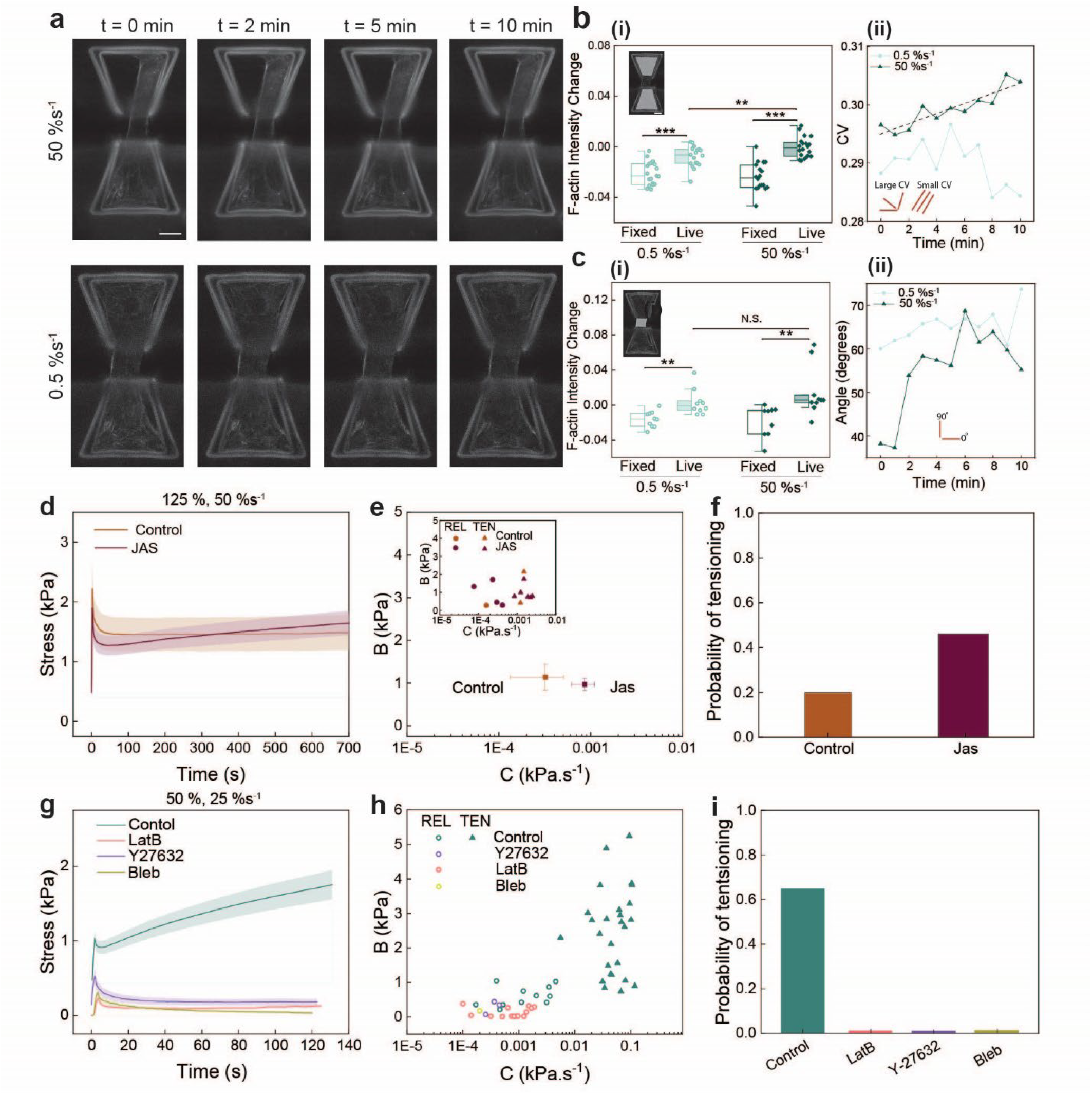
Actomyosin regulates the response of cell pairs in the ramp-and-hold experiments. **a.** Time series of F-actin in cells stretched to 50% strain at 50 %s^−1^ and 0.5 %s^−1^. Scale bar: 10 μm. **b**. Relative change in F-actin intensity (i) and circular variance (CV) of F-actin fibers over time (ii) while the strain was kept constant within the confinement region. Scale bar: 10 μm **c**. Relative change in F-actin intensity (i) and the angle of fibers relative to the stretching direction over time within the gap region (90° is parallel to stretching direction) near the cell-cell junction while the strain was held constant. Scale bar: 10 μm. **d**. The evolution of stress over time in control (N = 10, orange) and Jasplakinolide (Jas) treated (N = 13, brown) cell pairs stretched at 50 %s^−1^ to 125% strain (average of all curves for each condition ± s.e.). **e**. The average of the B vs C values calculated from fitting individual curves for the Jas treated cell pairs and the control condition on a semi logarithmic scale. Inset shows the clustered B vs C values where solid triangles and hollow circles represent tensioned and relaxed responses, respectively. **f**. The probability of tensioning for the control and Jas treated cell pairs in response to 125 % strain applied at 50 %s^−1^. **g**. The evolution of stress over time in control (N = 40, green), Bleb (N = 10, olive green), Y-27632 (N = 7, purple), and Lat B (N = 16, pink) treated cell pairs stretched at 25 %s^−1^ to 50% (average of all curves for each condition ± s.e.). **h**. The corresponding B vs C values calculated and clustered for the data presented in (**g**). **i**. The probability of tensioning for the cell pairs treated with actomyosin modulators and the control condition in response to 50% strain applied at 25 %s^−1^.

Next, we investigated the response of actin filament organization and orientation to strain. We first asked if strain could align F-actin by calculating the circular variance, which measures the dispersion of F-actin fibers (See Methods). Within the confinement region, CV was found to have a small but significant positive correlation with time for live cell pairs stretched at 50%*s*^−1^, but not at 0.5%*s*^−1^ (Fig. 4b (ii)). This suggests that only when cells experience the higher strain rate, F-actin fibers become more disorganized as the strain is held. Next, the change in orientation of the F-actin fibers relative to the stretching direction over time was measured within the gap region near the cell-cell junction (See Methods). Here, a small but significant positive correlation between orientation and time was found for live cells stretched at both rates (Fig. 4c (ii)). This again suggests that responses within this region are dominated by the stretching magnitude but not the rate.

One reason that cells may no longer be able to tension above 50% strain could be that the actin filaments rupture in response to stretch and prevent myosin mini filaments from applying tension. Therefore, we theorized that by stabilizing the actin cytoskeleton, tensioning could occur even at larger strains. To test this hypothesis, we repeated our experiments with cells treated with jasplakinolide (Jas), which stabilizes actin filaments. When we applied a strain of 125% at 25 %s^− 1^, Jas had a minimal influence on the magnitude of stress in cell pairs compared to controls (Fig. 4d). However, a tensioning behavior was qualitatively apparent in Jas treated cells. After analyzing individual curves, we found that the average C with Jas treatment is significantly higher than in the control group (Fig. 4e, Fig. 4e inset, and Supplementary Fig. 11b). Additionally, with Jas treatment, the probability of tensioning more than doubled, increasing from 20% to 46% (Fig. 4f), suggesting that, with a stable F-actin cytoskeleton, tensioning persists at large strain. These results point to the actin cytoskeleton as an important component in controlling the tensioning behavior of cell pairs.

We next asked if myosin contractility is required for tensioning. To test this idea, we examined the effect of drugs that destabilize the F-actin scaffold and inhibit myosin directly or indirectly. Tests were conducted at a strain of 50% at a rate of 25 %s^−1^, a condition shown to induce tensioning behavior in a large portion of cell pairs (Fig. 2d, Fig. 3d). Latrunculin B (LatB, which inhibits actin polymerization), Y27632 (which inhibits myosin contractility by inhibiting Rho-kinase), and blebbistatin (Bleb, which directly inhibits myosin contractility), all reduced the average magnitude of stress experienced by the cell pairs compared to controls, with the average value of B for all treatment conditions being significantly reduced (Fig. 4g, Supplementary Fig. 11e). When we categorized individual curves into tensioning and relaxing behaviors (Fig. 4h), we found that all three treatments resulted in eliminating the tensioning response and reducing the probability of tensioning to zero (Fig. 4i). Thus, myosin contractility and an intact F-actin network are essential components in the tensioning process.

### Tensioning is compatible with an active rheological model for actomyosin

We next sought to provide a rheological framework that encompasses all our observations. Based on our experiments, we posited the existence of a cellular detection mechanism responsible for triggering the tensioning of the cells that is sensitive to strain rate. Following detection, the signal is transduced into a response that increases tension at a constant rate through a mechanism that depends on actomyosin activities (Fig. 4). We therefore adapted a model of the active rheology of actomyosin (ARA) by Etienne et al.^29^ that accounts for both the contractility of myosin and turnover of the F-actin network to capture the dynamics of stress generation (Fig. 5a). Since we proved already that the parameters A and *α*, which control the short-term power-law relaxation, do not correlate with the probability of tensioning, we focused our attention on the longer-term response in relation to B and C. We assume that when deformation is applied, the detection mechanism elevates the steady state tension in cells to a new value, *σ*_*A*_. The ARA model uses this stress generation element in parallel to a dashpot, *η*_*A*_, and in series with an elastic component, *E*_1_. It provides us a simple way of testing this hypothesis and interpreting the rate of tensioning over time: upon a step change in contractile activity *σ*_*A*_, the system’s stress is expected to evolve towards a new plateau value at a rate controlled by both the compliance of the system and the turn-over of actomyosin (encompassed in the *η*_*A*_ term of the model). We could therefore interpret the growing tension as a transient but slow evolution of the system towards a new state of higher contractile tension. In the tensioning cases, where the stress can visibly curve towards a plateau, all parameters can be calculated; otherwise, in relaxed curves with no plateau where *η*_*A*_ does not converge, the mean of *η*_*A*_ from the tensioning cases is used to calculate *σ*_*A*_ and *E*_1_.

**Figure 5.**
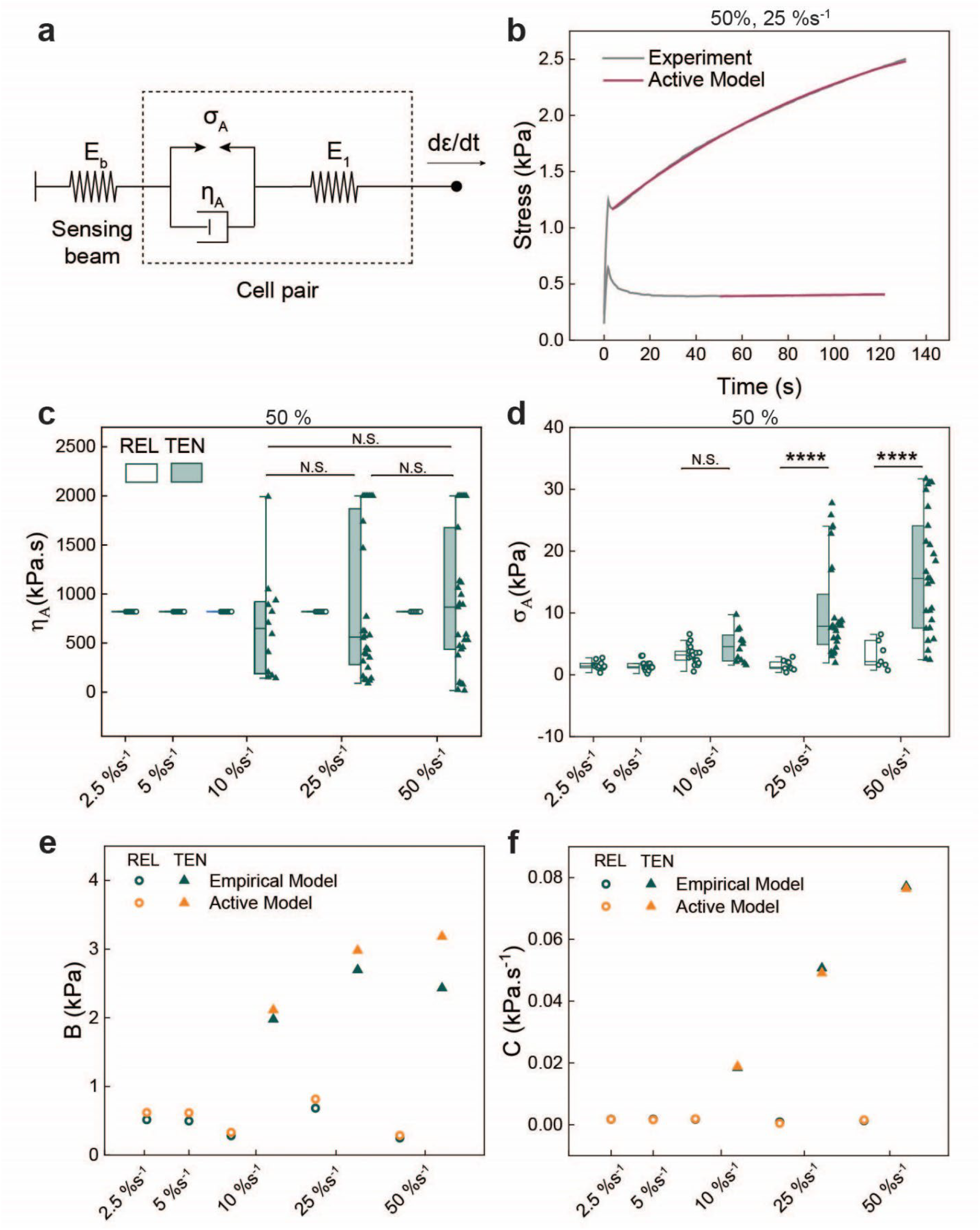
An active phenomenological model characterizes the bimodal stress response of the cell pair under different strain rates. **a.** A schematic illustration of the active phenomenological model which includes a spring labeled *E*_1_ symbolizing the elastic modulus of the cell pair. The model also incorporates an active branch, consisting of an active element *σ*_*A*_ to signify the contractile stress generation in the actomyosin network, along with a dashpot labeled *η*_*A*_, to represent the dissipation of the force generated by the myosin motor proteins through a viscous-like mechanism. **b**. The average relaxed and tensioned stress-time curves of the cell pairs stretched to 50% strain at 25 % s^−1^ are fitted using Eq. (6) according to the active model. **c**. The values of *η*_*A*_ are calculated for tensioned curves at high rates (i.e., 10 %s^−1^, 25 %s^−1^, and 50 %s^− 1^). These values are then averaged and kept as a constant for the relaxed curves. **d**. The values of *σ*_*A*_ obtained from fitting the individual curves for different rates with Eq. (6). **e, f**. Comparing the values of *B* (e) and *C* (f) derived from fitting the empirical and the active (*B* = *E*_*eq*_ and 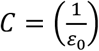 · 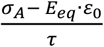, where 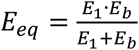 and 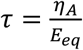) models on the average of the relaxed and tensioned curves for the data collected for 50% strain applied at various rates.

When we fitted our experimental data for 50% strain with the ARA model, we were able to accurately describe the prolonged response of the cell pairs in ramp and hold experiments and found good agreement between our model predictions and the data (Fig. 5b). We could also show that the ARA model parameters could be mapped to our B and C values in the transient linear phase (*B* = *E*_*eq*_ and 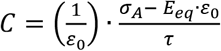, where 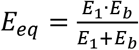 represents the spring equivalent of the cell pair and the sensing island and 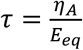 is the characteristic time determining the transition between the elastic-like to fluid-like response, see Methods, and Fig. 5e-f). When we examined the model parameters obtained by fitting individual curves, we found that the tension *σ*_*A*_ is significantly larger when the cell-pair is tensioning than when it is relaxing, whereas no obvious trend was apparent for *η*_*A*_ (Fig. 5c, d). Interestingly, the characteristic timescale of the contractile force generation 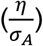 is on the order of 100 seconds, consistent with the reported actomyosin turnover timescale^5, 22^. This analysis shows that the steady tensioning process may result from a comparatively rapid but sustained change in the active stress state of the cell.

## DISCUSSION

Cells exhibit different behaviors in response to mechanical stimuli, including stress relaxation and reinforcement^5, 30, 31, 32, 33^. Stress relaxation is commonly observed in epithelial tissue and individual cells^5^. Another outcome resulting from mechanical stimulation is reinforcement, which can manifest as an increase in stiffness or larger force generation^31, 34, 35^. Our findings on epithelial cells reveal the existence of a new and different behavior in single cell pairs in response to strain.

Specifically, we identified an active tensioning response, where stress dynamically increases in response to ramp-and-hold strains at high rates resulting in higher stress than that arising passively from deformation. This response resembles the behavior of muscle fibers which stiffen in response to pulling at different speeds^36^ and it may serve as a protective mechanism physiologically to prevent tissue damage or fracture, by making further deformations difficult or even decreasing deformation (as shown in Supplementary Fig. 3b).

Our study shares similarities with the study by Khalilgharibi et al^5^, where they reported stress relaxation in MDCK epithelial monolayers and single cells with similar relaxation patterns as those we observed. Strikingly, in addition to stress relaxation, we observed that a significant proportion of cells increase the tension they generate over time. In contrast to the suspended anchorless monolayers which only exhibit relaxation, cell pairs and single cells in our study are fully anchored to a substrate via cell-ECM adhesions. These adhesions are known to participate in mechanosensing^37, 38, 39^, and our study points towards this adhesion site as the point of detection for cells to initiate their tensioning response. In addition to mechanosensing, a fully anchored cell can use its focal adhesion sites to physically balance the actomyosin generated tension in the actin network^40, 41^. Importantly, the tension response is sensitive to strain and strain rate. Our observations reveal clear functional limits to the tensioning response, as too high deformations or deformation rates lead to a muted or absent tensioning response.

Previous studies have also demonstrated that the rate of strain application plays a role in the level of force generation in fibroblast cells. Webster et al.^32^ have shown when a fibroblast cell is strained using a tipless AFM probe at low strain rates, force generation in the cell remains relatively unchanged compared to the force at the initial state. However, when the cell experiences higher strain rates, it exhibits increased contraction aiming to reach a new force set point and no stress relaxation. This behavior, termed tensional buffering, allows cells to preserve their structural integrity by adjusting to the intensity of mechanical stimuli which can be slow in processes like tissue development or rapid due to injury-related forces. Additionally, they also report a limit for the increased contraction level, with cells displaying stress relaxation when they undergo a step strain, similar to our findings. Whether this threshold is due to limitations in sensing or in transduction remains unclear. Force transmission to cell-ECM adhesions could be disrupted due to rupture at cell-cell junctions under high deformations. Preliminary results show E-cadherin reorganization at the junction in response to the elevated stress at high rates in the functional range and point towards its role in maintaining the mechanical link to facilitate sensing at the cell-ECM adhesion (Supplementary Movie 4). In all cases, tensioning arose within ~10s of application of deformation. Such a time-scale points to mechanisms involving signalling in the form of post translational modifications, ion channels, or emergent mechanisms such as the catch bond properties of myosin^42^.

We speculate that deformation applied at different rates could induce sensing at cell-ECM adhesions via a mechanism similar to substrate stiffness detection. This may be facilitated by force-induced unfolding of proteins such as talin, followed by vinculin recruitment^7, 43, 44, 45^. Under actomyosin generated tension, protein unfolding and subsequent mechanosensing is induced only for stiff ECM, which allows sufficient force build-up^45^. Under applied strain, tension is applied to this same adhesion complex through stretch of the basal side of the cell and activates potentially the same detection mechanism. This activation occurs during the ramp strain application and influences cell behavior during holding. More importantly, studies have shown that detection molecules are not only sensitive to the magnitude of the force, but also the rate at which it is applied and this can be described by the interplay between the rheology of the cells and the molecular clutch model^7^. Our observation of higher tensioning probability at high strain rates also agrees with this model, in which high rates (50 %s^−1^ compares to 40 %s^−1^ in Andreu et al.^7^) allow faster force build-up in sensing molecules, and thus higher probability of triggering a response. The exact conditions at the cell-ECM and cell-cell adhesion sites, such as force-sensing protein distribution and degree of clustering, will influence how stress is distributed across these proteins and can influence the likelihood of triggering the range of responses. Future research using FRET sensors integrated within mechanosensory proteins (e.g. talin and vinculin) will be necessary to explore the molecular mechanism triggering tensioning. The higher propensity of stress relaxation at large strain may be a sign of actin network disruption or damage due to the strain application, resulting in cells failing to perform force generation in response to high strain rates even when stimuli is detected^7, 27, 32^. Consistent with this idea, when we applied large strain in two steps separated by a pause rather than a single step, the tensioning response was preserved, suggesting that the pause allows the cell to adapt, perhaps through remodeling of the cytoskeleton. Damage to the actomyosin cytoskeleton could arise because of loss of connectivity within the actin network^46, 47^, similar to the loss of contact between myosin thick filaments and F-actin thin filaments when sarcomeres are overextended^48, 49^. Alternatively, damage could arise from direct breakages in actin filaments due to the large and rapid application of strain^7, 20, 50^. In support of this, stabilization of F-actin with Jas enabled tensioning responses to persist at high strain. However, further work will be necessary to determine the mechanism underlying loss of tensioning.

## METHODS

### TPP fabrication

Fabrication of stretchers was carried out as described in ref.^8^. Briefly, standard tessellation language (STL) files were generated from the geometry of the stretcher made with COMSOL 4.2 software and were imported into the Describe software (Nanoscribe, GmbH) to compile job files for the Photonic Professional (GT) tool (Nanoscribe, GmbH). The stretchers were fabricated from IP-S photoresist (Nanoscribe) with a 25× immersion microscope objective. Glass coverslips with diameters between 11 and 25 mm and thickness of ~160 μm were coated in indium tin oxide (ITO) to allow autofocusing. To promote adhesion of the stretchers to the glass slide, a ~2 μm thick layer of porous silicon oxide (PSO) was deposited onto the ITO-coated slides using an in-house developed protocol. Arrays of stretchers varying from 5×5 to 6×6 were fabricated on each slide.

### Cell culture

A431 cells were cultured in growth medium composed of Dulbecco’s modified Eagle’s medium (ThermoFisher) supplemented with 10% FBS and 1% P/S in an incubator at 37°C with 5% CO_2_.

### Stretcher preparation and cell deposition

The glass slide on which the stretchers were fabricated was adhered to a glass bottom petri dish using an optically clear UV curable adhesive (Norland Products). To prepare stretchers for cell deposition, a 100 μL drop of Geltrex (Thermofisher) was placed on the stretchers and incubated for 1 hour to promote adhesion of cells to the stretchers. Finally, the remaining Geltrex solution was rinsed off and the petri dish was filled with CO_2_ independent media (Thermofisher) supplemented with 10% FBS, 1% P/S, and 1% GlutaMAX (Thermofisher). Cells were deposited on the stretching platforms using the protocol described in ref.^8^. Briefly, an Eppendorf single-cell isolation setup was used to pick up and place cells within the confinement on the platform. The setup consisted of a microcapillary (Piezo Drill Tip ICSI, Eppendorf) connected to a pressure controller (CellTram 4r Air/Oil, Eppendorf), which was mounted on a three-axis micromanipulator (Transferman 4R, Eppendorf). After a drop of cells was placed within the petri dish containing the stretchers, cells were picked up and placed on the stretchers one at a time. Cell deposition was carried out on a Nikon Eclipse Ti-S microscope fitted with a custom heating chamber maintained at 37°C. After cells were deposited the petri dish was kept on the microscope overnight to allow cells to adhere to the stretchers and form cell-cell adhesions with adjacent cells on the same stretchers.

### Mechanical stretching test

Cell pairs were subjected to mechanical stretch using an AFM as described previously^8^. Briefly, a through hole was drilled into the end of an AFM cantilever tip (NanoAndMore) with a focused ion beam to engage with the pillar on the forcing island. With the hole engaged with the pillar, the displacement and displacement rate were selected in the AFM software and stretch was initiated. The tests were visualized with a Zeiss Axio Observer 7 microscope and recorded using a screen-recording software (Camtasia). After stretching experiments were complete, the petri dish was incubated in trypsin for 1 hour to detach cells from the stretchers. The stretchers were rinsed with PBS to remove cells, and finally submerged in 70% ethanol for 1 minute to sterilize the stretchers in preparation for the next experiment. Stretchers could be reused up to 10 times before they began detaching from the substrate, at which point new stretchers were fabricated.

### Drug treatment

For conducting cell stretching experiments with drug treatments, cells were treated with the drug immediately before stretching them. For latrunculin B (Abcam) treatment, LatB was diluted in DMSO and then diluted in DMEM to a final concentration of 200 nM and exposed to the cells for 30 minutes. For blebbistatin (Sigma Aldrich) treatment, stock bleb was diluted in DMSO and then diluted in DMEM to final concentrations of 3.4 and 4.25 μM and exposed to cells for 2 hours. For Y27632 (Abcam) treatment, stock Y27632 was diluted in DMSO and then diluted in DMEM to a final concentration of 30 μM and exposed to cells for 10 minutes. For Jasplakinolide (Thermofisher) treatment, stock Jasplakinolide was diluted in DMSO and then diluted in DMEM to a final concentration of 0.1 μM and exposed cells for 10 minutes. For all drug treatments the media containing the drug was replaced with fresh media before beginning the stretching experiment.

### Displacement tracking and stress-time curve calculation

The displacement of each island and the calculation of stress were carried out as described previously^8^. Briefly, the location of each island was determined using a modified version of MATLAB DIC. A rectangular region was defined along the confinement bowtie on each island and the location of each pixel within this region was tracked on every tenth frame to reduce computation time. The displacement of each pixel relative to the first analyzed frame was determined and then the average of the displacement of each pixel was calculated for each island to yield the displacement of each island. The MATLAB code is available at github.com/YangLabUNL.

To calculate stress, first the force within the cell pair system was determined based on the displacement of the sensing island and Hooke’s Law, *F* = *kδ*, where *k* is the previously calculated stiffness of the bottom island, 0.11 N/m. Engineering stress was calculated based on the measured force and the original cross-sectional area of the cell-cell contact, which was determined to be 120 *μm*^2^ with little variation due to the geometry of the confinement. This value was obtained by multiplying the length and the thickness of the intercellular junction based on fluorescent images^8^. Finally, the time for each data point was determined based on the framerate of the original video and the percentage of analyzed frames to give the stress-time curve.

To calculate strain, the deformation of the cell pair or single cell was divided by its initial length, following the equation *ε* = *D*/*L*_0_, with the deflection of the sensing island δ assumed to be negligible compared to the forcing island displacement. For a cell pair, the initial length was 20 μm, considering the nucleus-to-nucleus spacing determined previously^8^. As shown in Supplementary Fig. 1, the nucleus-to-nucleus region experiences noticeable deformation during the loading phase, whereas the areas beyond the nucleus remain relatively unaffected. and for a single cell was 17 μm, or half the length of the confinement along the stretching direction.

### Tracking of F-actin intensity

After cell deposition, cells were treated with SPY650-FastAct (Cytoskeleton) at a 1X concentration for 2 hours prior to imaging. For control experiments, cells were fixed with 4% paraformaldehyde for 10 minutes prior to imaging. Imaging was carried out on a Ziess LSM 800 confocal microscope, and images of the cell pairs were taken once a minute. Intensity change was quantified by determining the average intensity within the given region of interest (either within the confinement or between the islands) at each timepoint and calculating the slope of the line of best fit. Therefore, positive values represent an increase in intensity and negative values represent a decrease in intensity.

### Dispersion and angle of F-actin

Each image was divided into 16 × 16-pixel grids, and for each grid, the dominant fiber orientation was calculated using Fourier Transform (FT) analysis. This technique converts the image from the spatial domain to the frequency domain, enabling the extraction of key structural features, such as fiber orientation and anisotropy level. From orientation data, two parameters were calculated for each image. The first parameter, circular variance (CV), quantifies the dispersion of fibers within the two-dimensional imaging plane. CV values range from 0, indicating perfectly aligned fibers, to 1, indicating complete dispersion. A modified definition of CV was used to tailor the analysis specifically for fibrous structures (see reference ^51^) The second parameter, the average angle (Angle), represents the orientation of fibers relative to an axis perpendicular to the stretching direction. An Angle of 0° indicates fibers oriented perpendicular to the stretching direction, while an Angle of 90° indicates a parallel orientation. Calculations included only those grids classified as anisotropic by the FT analysis. For more details see [reference ^52^].

### Data analysis and modeling

The stretching process applied to our cell pairs involved two phases. Initially, a stretch was applied at a constant rate to reach the desired strain magnitude *ε*_0_, followed by keeping the strain constant for a period of two minutes. To account for the impact of the rate of strain application during the initial phase, we utilized a convolution integral to accurately model the temporal evolution of stress *σ*(*t*) in our experiments:

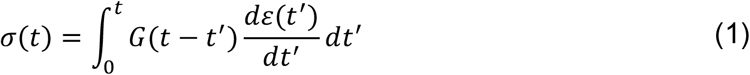

In our model, we define the relaxation modulus 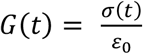 as *G*(*t*) = *At*^−*α*^ + *B* + *Ct*. This equation incorporates several terms: a power law term *At*^−*α*^ to capture the relaxation phase occurring within the first few seconds, *B* · *ε*_0_ representing the residual stress in the relaxed curves or an estimate of the point at which stress starts to increase in the tensioned curves, and finally, the linear function *Ct* to account for the stress increase in the tensioned curves. We employed the RHEOS software implemented in the Julia programming language to fit our experimental data^53^.

To initiate the fitting procedure, we assign initial values to each parameter of the *G*(*t*) function. The initial value of *B* is defined as the lowest stress level occurring between *t*_*hold*_, which represents the second time point after maintaining the applied stretch constant, and the final point at the end of the two-minute hold (Supplementary Fig. 3a). The initial value for *α* is calculated as the slope of a line fitted on the (*σ*(*t*_*hold*_ < *t* < *t*_*σmin*_) − *B*) on a log-log scale (Supplementary Fig. 3c-d). Here, *t*_*σmin*_ refers to the time at which the initial value for B was calculated. Then, the curves are cut at *σ*_*cut*_ = 1.4_*σmin*_, and the slope of the line fitted on the *σ*(*t*_*σmin*_ < *t* < *t*_*σcut*_) is defined as the initial value for C (Supplementary Fig. 3f). Finally, the initial value for A is obtained by fitting the curve with the function *σ*(*t*) = *A*^′^*t*^−*α*^ + *B*^′^ + *C*^′^*t*, where the values of *α, B*^′^, and *C*^′^ are the initial values calculated in the previous steps. To ensure the consistency of the outcomes for the fittings, we employed three optimization methods, namely COBYLA, BOBYQA, and SBPLX. These techniques yielded comparable errors in most cases (Supplementary Fig. 3g-h). The fitting parameters presented in this study are derived from the fitting process conducted with the BOBYQA optimizer.

Given the observation of two visually distinguishable behaviors in the experimental data presented in Fig. 2a and Supplementary Fig.2, we aimed to identify a mathematical approach to categorize the data. To accomplish this, we initially plotted the values of B vs C on a semi-logarithmic scale (Fig. 2d), which highlighted the existence of two distinct groups. Subsequently, we employed the NbClust package to determine the optimal number of clusters for our dataset, and the analysis confirmed that two clusters were the most appropriate choice. In the next step, to eliminate bias in the results of our clustering, we applied two clustering algorithms, namely K-means and K-medoids, to classify the dataset into two groups. The outcomes from both clustering algorithms exhibited similarity, effectively categorizing the experimental results into two groups: relaxed and tensioned (Supplementary Fig. 4).

### Statistics and reproducibility

The excluded outliers were defined as the values outside of the range [*Q*_1_ − 1.5(*Q*_3_ − *Q*_1_), *Q*_3_ + 1.5(*Q*_3_ − *Q*_1_)], where *Q*_1_ and *Q*_3_ are the 25th and 75th percentile, respectively. The normality of each dataset was tested using Shapiro-Wilk tests, and equality of variances was tested using F-tests. When the data was drawn from a normally distributed population, the selection of the analysis method depended on the equality of variances among different data groups. When examining two groups of data, we conducted an independent sample t-test in cases of equal variances, and for cases of unequal variances, we employed Modified t-tests with Welch correction. For groups consisting of three or more, One-way ANOVA was utilized when variances were equal, while Welch’s ANOVA was applied in instances of unequal variances. Subsequently, to identify statistically significant differences among group means, we conducted Tukey’s post hoc tests for groups with equal variances and Games Howell’s post hoc tests for groups with unequal variances.

If the data did not exhibit a normal distribution, we employed the Mann-Whitney test for the analysis of two data groups and the Kruskal-Wallis nonparametric ANOVA test with Dunn’s post hoc test for three or more data groups. We conducted statistical analysis using the OriginLab software. Datasets with *p* < 0.05 were considered to have significantly different means and are denoted by a single asterisk (*). Data sets with *p* < 0.01, *p* < 0.001, and *p* < 0.0001 were highly significantly different and were denoted by double (**), triple (***) and quadruple (****) asterisks, respectively. The edges of the box plots represent the 25th and 75th percentiles of the data, the line in the middle marks the median, and whiskers extend to include maximum and minimum values in the dataset. Experiments for each group of data were repeated at least three times to ensure reproducibility. The number of samples are included in the figure captions.

### Active phenomenological model

Here, we employed an active phenomenological model (Fig 6a), which has been previously utilized by Etienne et al.^29^ to explain how single cells respond to changes in the stiffness of their environment. Our goal was to apply this model to characterize the strain rate dependent response of cell pairs to a sustained strain. This model comprises two main components: a spring represented by *E*_1_, signifying the elastic modulus of the cell pair, and an active portion that includes an active element denoted as *σ*_*A*_, representing the contractile stress generated by actomyosin activity, in parallel with a dashpot labeled as *η*_*A*_, which accounts for the dissipation of force generated by myosin motor proteins through a mechanism resembling viscosity. Using this model, we were able to accurately describe the prolonged response of cell pairs in ramp and hold experiments.

The sensing island of the microstructure is modeled as a spring *E*_*b*_ in series with the model of the cell pair. These two springs (*E*_1_ and *E*_*b*_) are combined into a single equivalent spring of 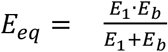. Stress and strain in the active section and the equivalent spring are presented below:

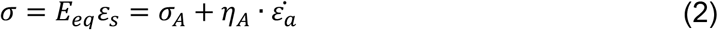

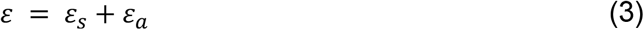

where *ε*_*s*_ and *ε*_*a*_ are displacements in the equivalent spring and the active section, respectively. Constitutive equation of the model is therefore the following equation:

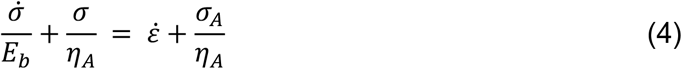

And the response of the model to a step strain *ε*_0_ is:

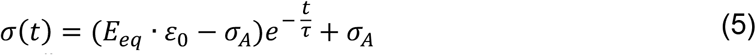

with the characteristic time scale 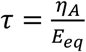. The relaxation modulus of the system is therefore:

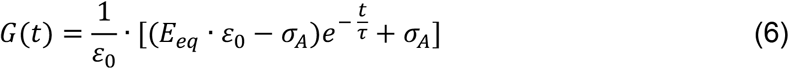

This relaxation modulus was then used in Equation (1) to fit the average relaxed and tensioned curves of the experimental data collected at 50% strain using the RHEOS software (Fig. 6b).

We have also shown that when *t* << *τ*, stress in the system can be defined as:

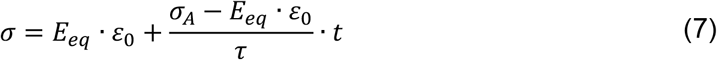

which is equivalent to the empirical model we presented before, *σ*(*t*) = *B* + *Ct*, with *B* = *E*_*eq*_ and 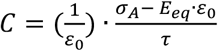 ·

## Data Availability

Source data for the presented graphs are provided in the Supplementary Data files. All other data supporting the findings of this study are available upon reasonable request.

## Code Availability

Codes for the modeling and clustering are available at GitHub: YangLabMSU.

## Acknowledgement

We acknowledge fundings from the NSF (award #1826135, #2143997), and the NIH National Institutes of General Medical Sciences (R35GM150623).

## Author contributions

B.T.S., J.R. and A.M.E. performed stretch-and-hold tests on cell pairs and single cells. J.R. and A.M.E. analyzed the tensile tests and produced stress-time and strain-time curves. G.M. conducted AFM characterization of the TPP structure for calibrating its stiffness. B.T.S. and A.K. developed an empirical function to analyze the stress-time curves. B.T.S., A.K. and C.H. utilized an active phenomenological model for data analysis. J. R. performed F-actin and E-cadherin imaging and analyzed them for their intensity. A.O.M. analyzed the F-actin images for dispersion and angle. N.V.L. fabricated the microstructures used for the tensile test. R.Y. conceived the project and designed the experiments. B.T.S., J.R., G.C., A.K., and R.Y. wrote the manuscript. All authors discussed the results and the manuscript.

