## Supplementary material for "Sustained Strain Applied at High Rates Drives Dynamic Tensioning in Epithelial Cells": SI

### Table of Content

Supplementary figures (1-13)

Supplementary table S1 (14)

Supplementary movie captions (15)

**SUPPLEMENTARY FIGURES**

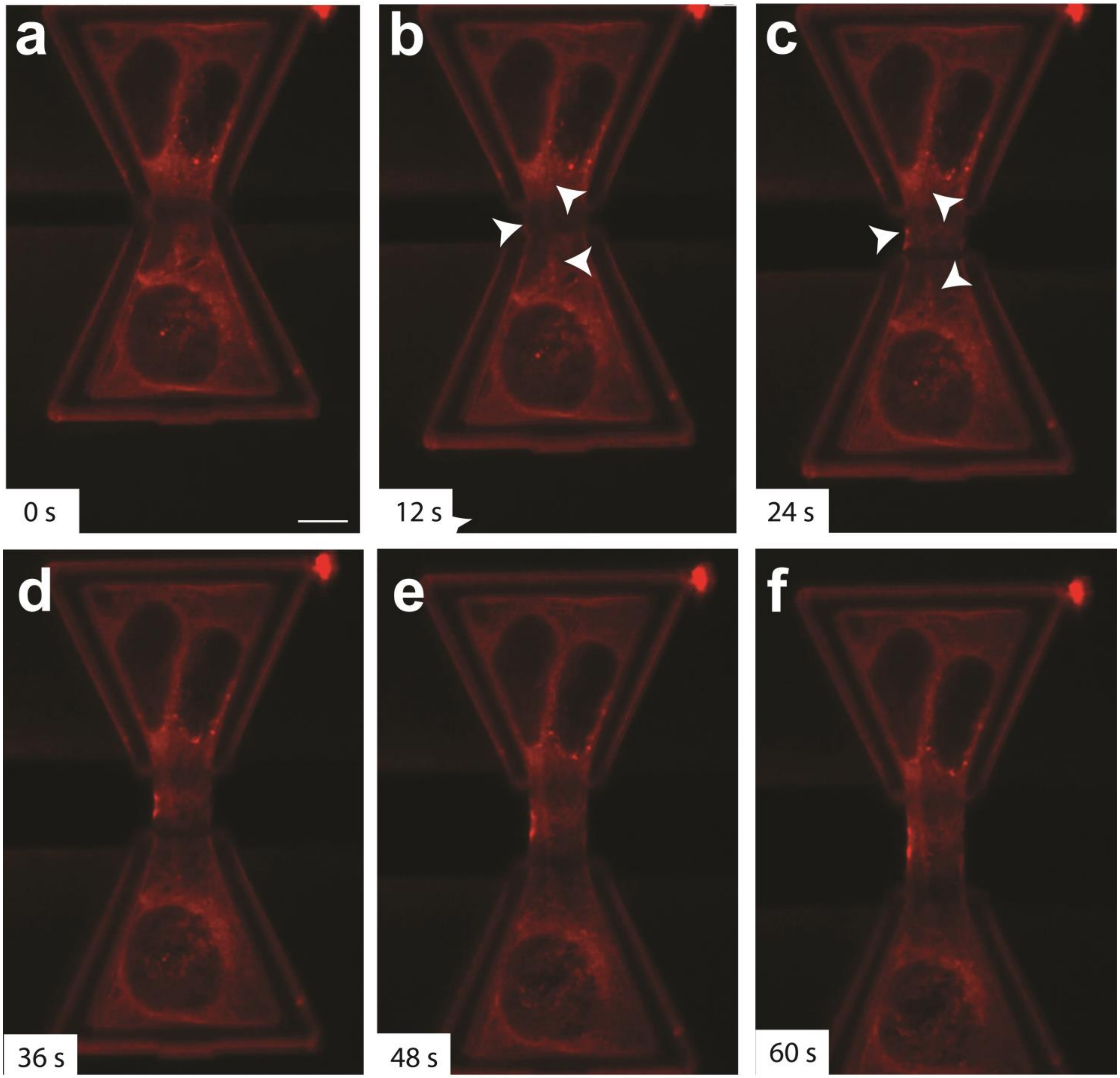


**Supplementary Fig. 1. Fluorescence images of actin cytoskeleton during the loading phase. a-f.** Representative images captured in 12 second intervals from an example stretch test at a rate of $0.55\%s^{-1}$ to highlight the deformation of the region between the cell nuclei. Scale bar: 10 µm.

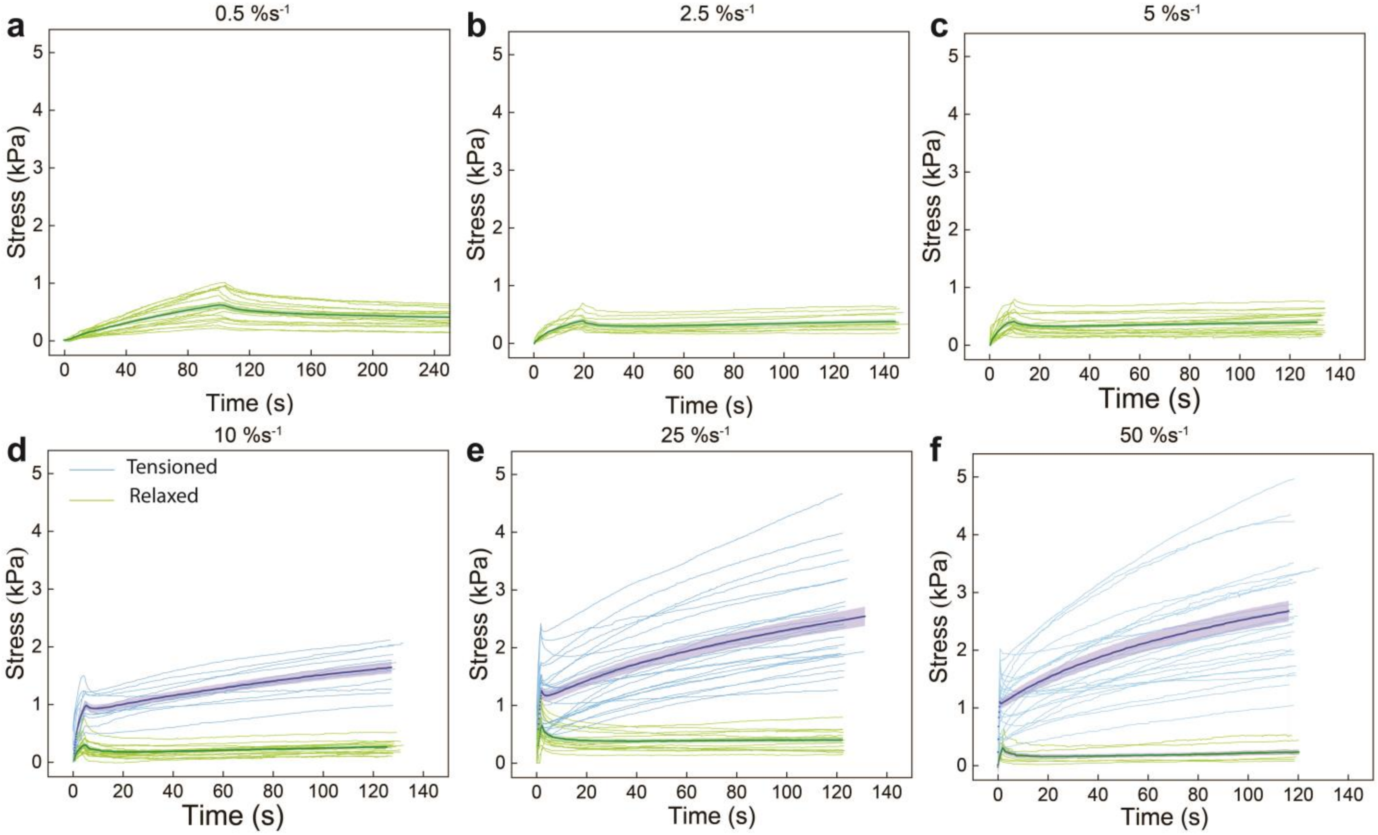


**Supplementary Fig. 2. The response of epithelial cell pairs to 50% strain applied at different strain rates. a-c.** Individual and average stress-time curves for the cell pairs stretched at 0.5 $\%s^{-1}$ (N=19) (**a**), 2.5 $\%s^{-1}$ (N=12) (**b**), and 5 $\%s^{-1}$ (N=21) (**c**). Based on the results of clustering of the B vs C data, we used the color green to demonstrate relaxation. **d-f.** Individual and the average stress-time curves for the cell pairs stretched at 10 $\%s^{-1}$ (N=30) (**d**), 25 $\%s^{-1}$ (N=40) (**e**), and 50 $\%s^{-1}$ (N=31) (**f**). At these strain rates the cell pairs display both relaxed and tensioned behaviors, as denoted by the colors green and blue, respectively.

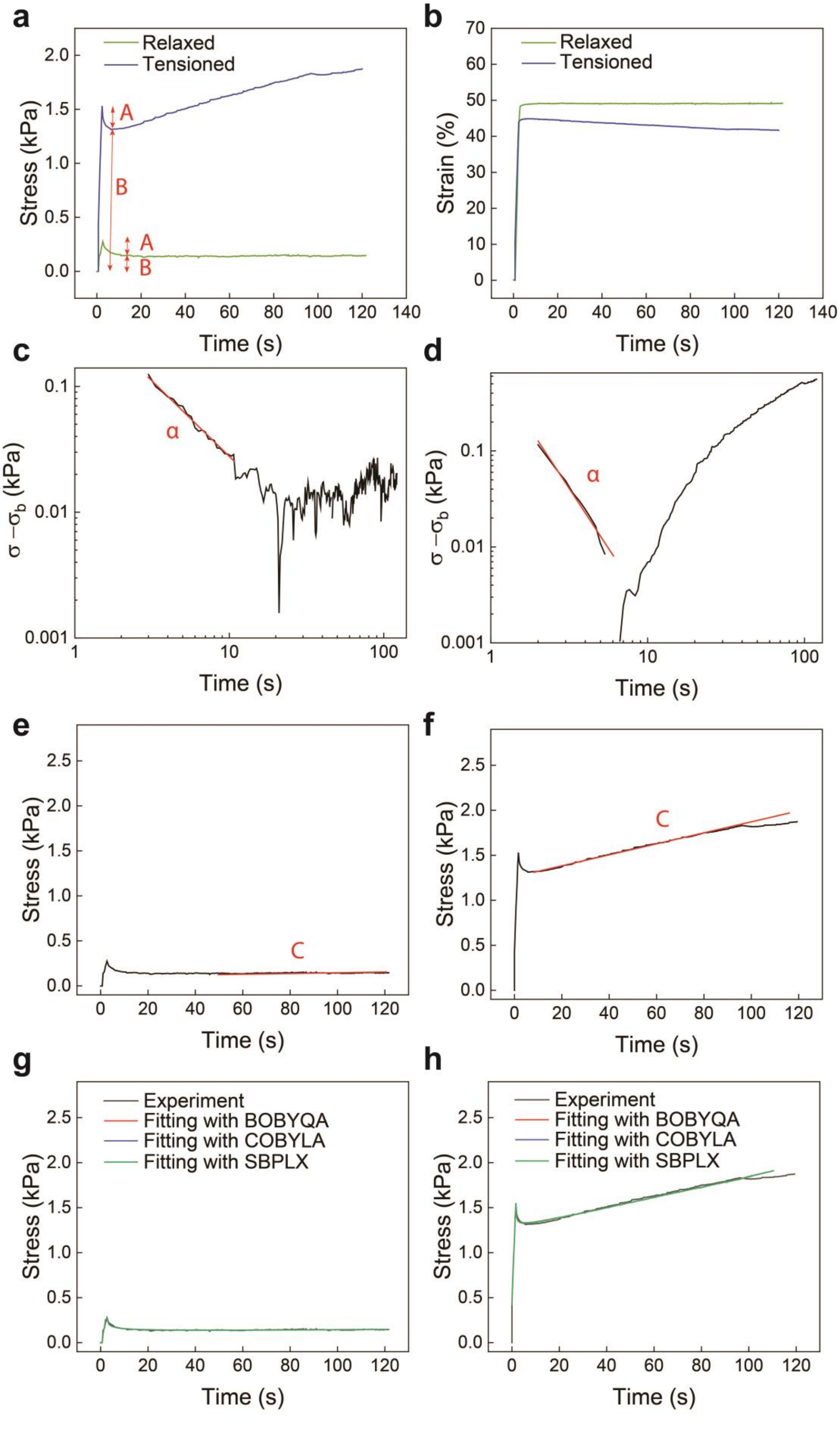
a
Relaxed
Tensioned
A
B
A
B
Stress (kPa)
Time (s)
b
Relaxed
Tensioned
Strain (%)
Time (s)
c
α
σ −σb (kPa)
Time (s)
d
α
σ −σb (kPa)
Time (s)
e
C
Stress (kPa)
Time (s)
f
C
Stress (kPa)
Time (s)
g
Experiment
Fitting with BOBYQA
Fitting with COBYLA
Fitting with SBPLX
Stress (kPa)
Time (s)
h
Experiment
Fitting with BOBYQA
Fitting with COBYLA
Fitting with SBPLX
Stress (kPa)
Time (s)

**Supplementary Fig. 3. Characterization of the experimental results.** The function, $G(t) = At^{-\alpha} + B + Ct,$ is used for fitting the stress-time curves. **a, b.** Example relaxed and tensioned stress-time (**a**) and strain-time (**b**) response of cell pairs to 50% strain applied at 25 $\%s^{-1}$. $A \cdot \varepsilon_0$ is the amplitude of stress relaxation and $B \cdot \varepsilon_0$ is the residual stress. **c, d.** $\alpha$ is the slope of the stress-time curves on a log-log scale after removing the residual stress from the response in relaxed (**c**) and tensioned curves (**d**). **e, f.** $C \cdot \varepsilon_0$ is the slope of the stress-time curve from the point where the stress is minimum after holding the applied displacement constant to the end of the two minutes hold. **g, h.** Comparison between the results of the three optimization methods (BOBYQA, COBYLA, and SBPLX) that are used for fitting both relaxed (**g**) and tensioned (**h**) curves. The values of model parameters for the relaxed curve (**g**) are ($A$ = 0.242 kPa. $s^{\alpha}$, $\alpha$ = 0.43, $B$ = 0.214 kPa, $C$ = 0.00045 kPa.$s^{-1}$) and the model parameters for the tensioned curve (**h**) are ($A$ = 0.324 kPa. $s^{\alpha}$, $\alpha$ = 0.663, $B$ = 3.018 kPa, $C$ = 0.017 kPa.$s^{-1}$).

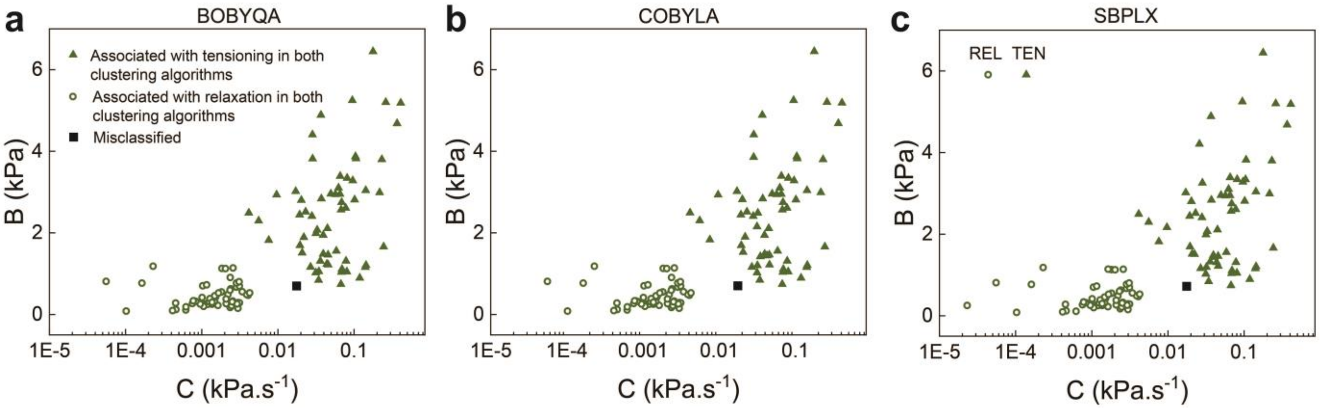


**Supplementary Fig. 4. Comparison between the results of the clustering algorithms. a-c.** The values of B vs C obtained from fitting the experimental results using three optimization methods, namely BOBYQA (**a**), COBYLA (**b**), and SBPLX (**c**), are plotted on a semi logarithmic scale. Then, data was clustered using two clustering algorithms: K-means and K-medoids. The points associated with tensioning in both clustering algorithms are shown with a green triangle, and the points associated with relaxation are represented with green hollow circles. Data points identified as tensioned in one algorithm and as relaxed in the other are illustrated as black squares.

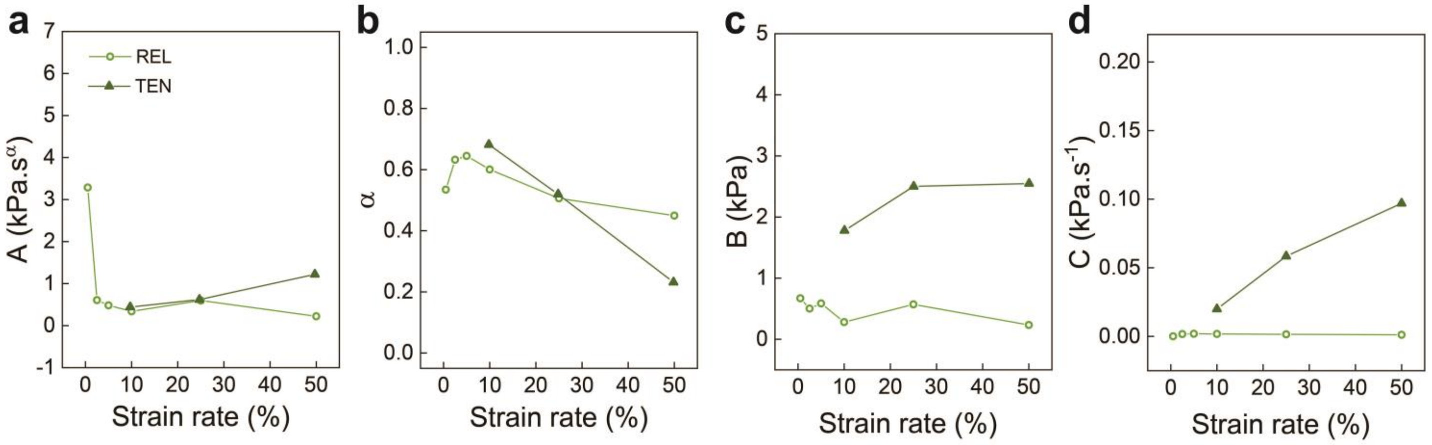


**Supplementary Fig. 5. Averages of the model parameters calculated for relaxed and tensioned curves at different strain rates to reach a 50% strain.** The model parameters A (**a**), α (**b**), B (**c**), and C (**d**) are plotted as a function of strain rate for relaxed and tensioned curves.

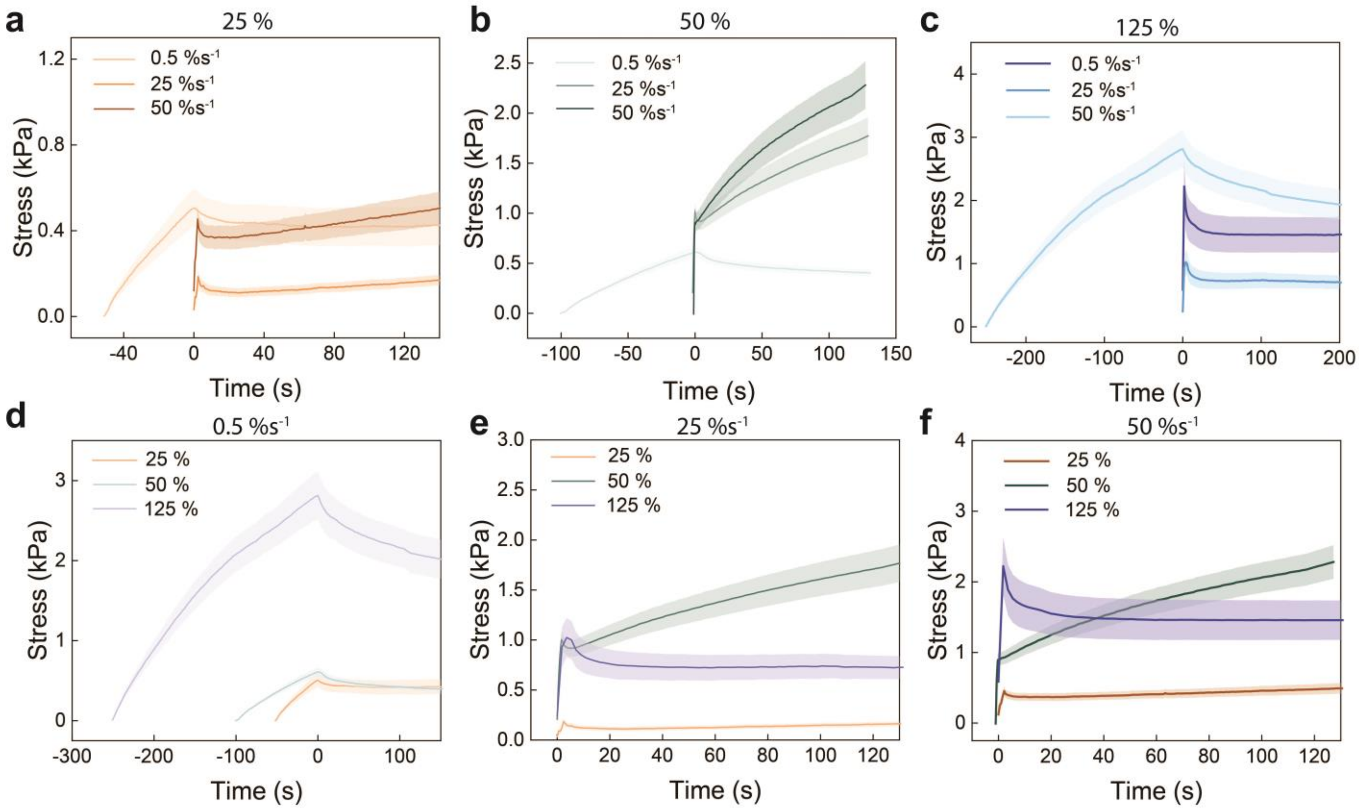


**Supplementary Fig. 6. Response of cell pairs to different magnitudes and rates of strain. a-c.** Temporal evolution of stress for the cell pairs following strains of 25% (**a**), 50% (**b**), and 125% (**c**) that are applied at three different rates. **d-f.** Response of cell pairs to different strain magnitudes at a constant rate of 0.5 $\%s^{-1}$ (**d**), 25 $\%s^{-1}$ (**e**), and 50 $\%s^{-1}$ (**f**). Each curve represents the average of all curves for that condition $\pm$ s.e.

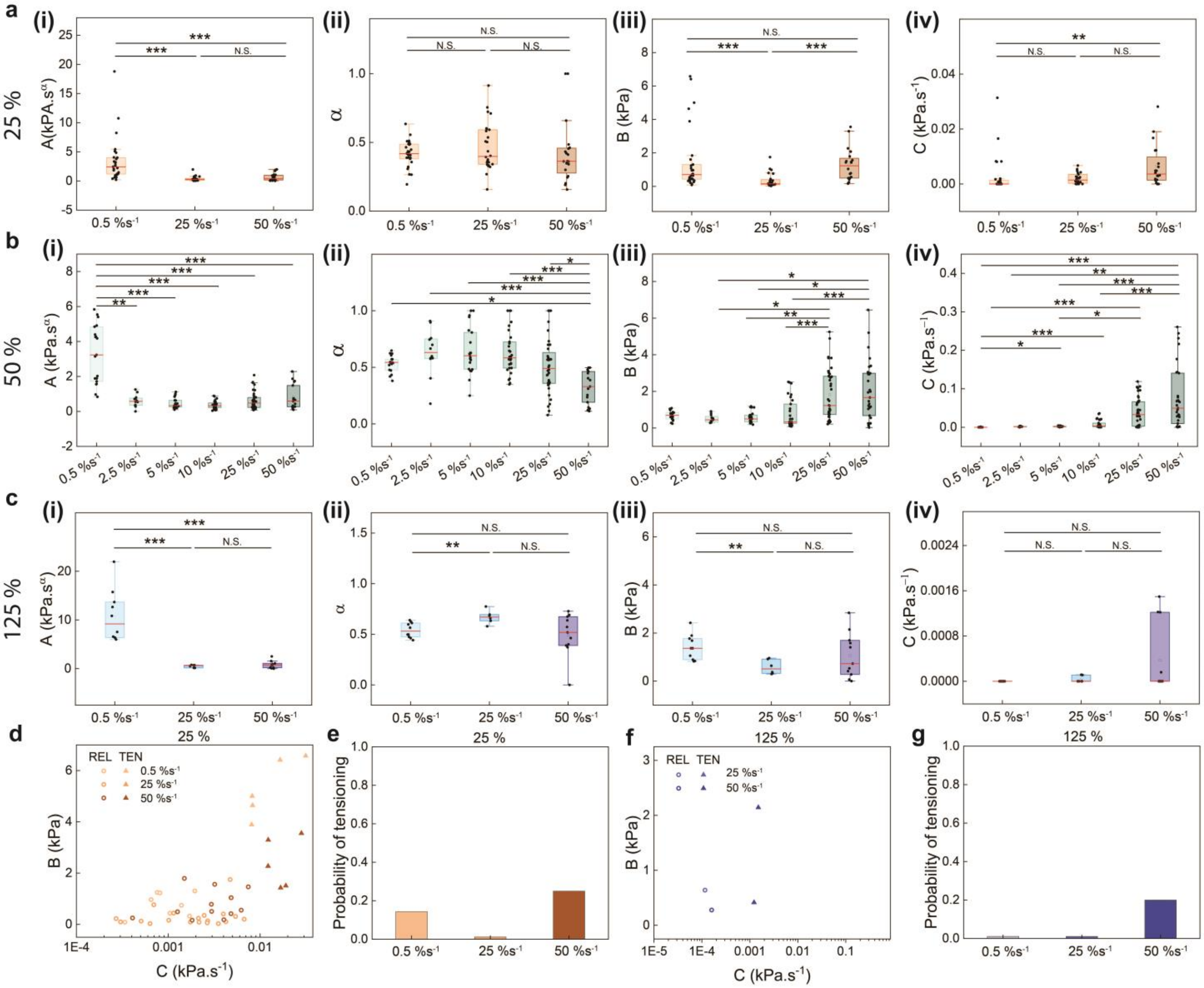

**Supplementary Fig. 7. Modeling parameters for studying the role of the applied strain rate. a-c.** Box plots representing parameters $A \cdot \varepsilon_0$ (amplitude of relaxation), $\alpha$ (power law exponent), $B \cdot \varepsilon_0$ (residual stress), and $C \cdot \varepsilon_0$ (captures the slope of the tensioning) for the cell pairs experiencing 25% (**a**), 50% (**b**), and 125% (**c**) strain for various levels of strain rate. **d**. B vs C values for all strain rates applied to reach 25% strain are plotted and clustered on a semi logarithmic scale. The results of clustering are represented by solid triangles and hollow circles to represent tensioned and relaxed curves, respectively. **e.** Probability of tensioning for different rates that are applied to reach 25% strain. **f.** B vs C values for all strain rates applied to reach 125% strain are plotted and clustered on a semi logarithmic scale. **g.** Probability of tensioning for different rates that are applied to reach 125% strain.

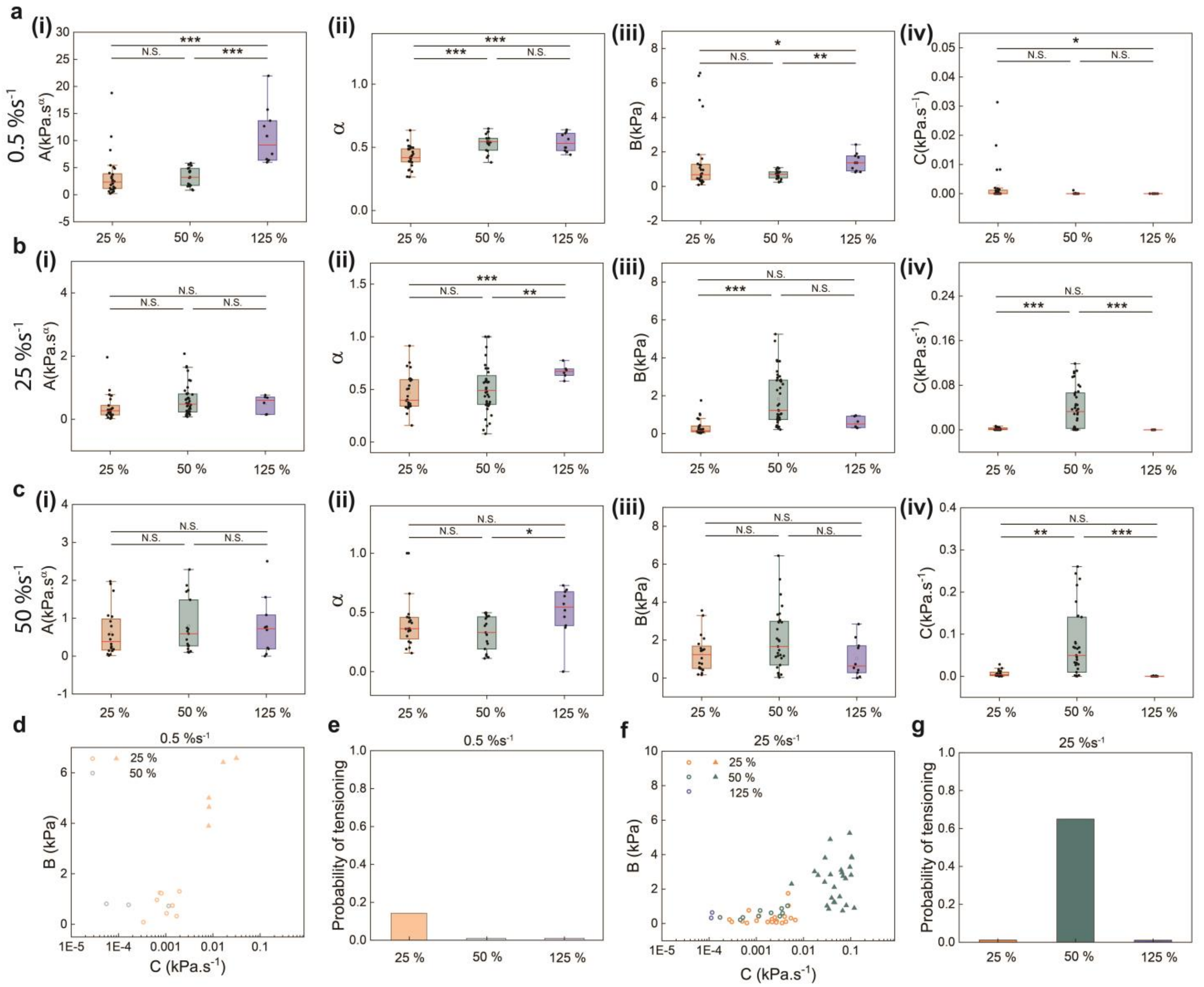


**Supplementary Fig. 8. Modeling parameters for studying the role of the applied strain magnitude. a-c.** Box plots representing parameters $A \cdot \varepsilon_0$ (amplitude of relaxation), $\alpha$ (power law exponent), $B \cdot \varepsilon_0$ (residual stress), and $C \cdot \varepsilon_0$ (captures tensioning) for the cell pairs experiencing different strain magnitudes at a constant rate of 0.5 %s$^{-1}$ (**a**), 25 %s$^{-1}$ (**b**), and 50 %s$^{-1}$ (**c**). **d**. B vs C values calculated from fitting the response of cell pairs to different strains applied at a constant strain of 0.5 %s$^{-1}$. The results of clustering are represented by solid triangles and hollow circles to represent tensioned and relaxed curves, respectively. **e.** Probability of tensioning for different strain magnitudes that are applied at a constant rate of 0.5 %s$^{-1}$. **f.** B vs C values calculated from fitting the response of cell pairs to different strains applied at a constant strain of 25 %s$^{-1}$. **g.** Probability of tensioning for different strain magnitudes that are applied at a constant rate of 25 %s$^{-1}$.

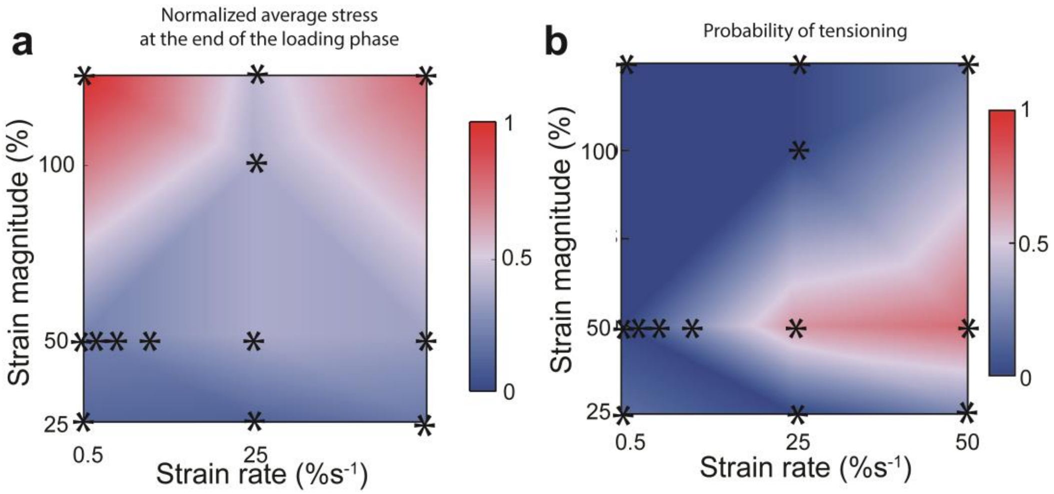


**Supplementary Fig. 9. Plots illustrate the normalized average values of stress at the end of the loading phase (a) and probability of tensioning (b) versus the strain magnitude and rate.** Stars represent values derived from experimental results. Higher values are indicated by the number 1 and the color red, while lower values are represented by the number 0 and the color blue.

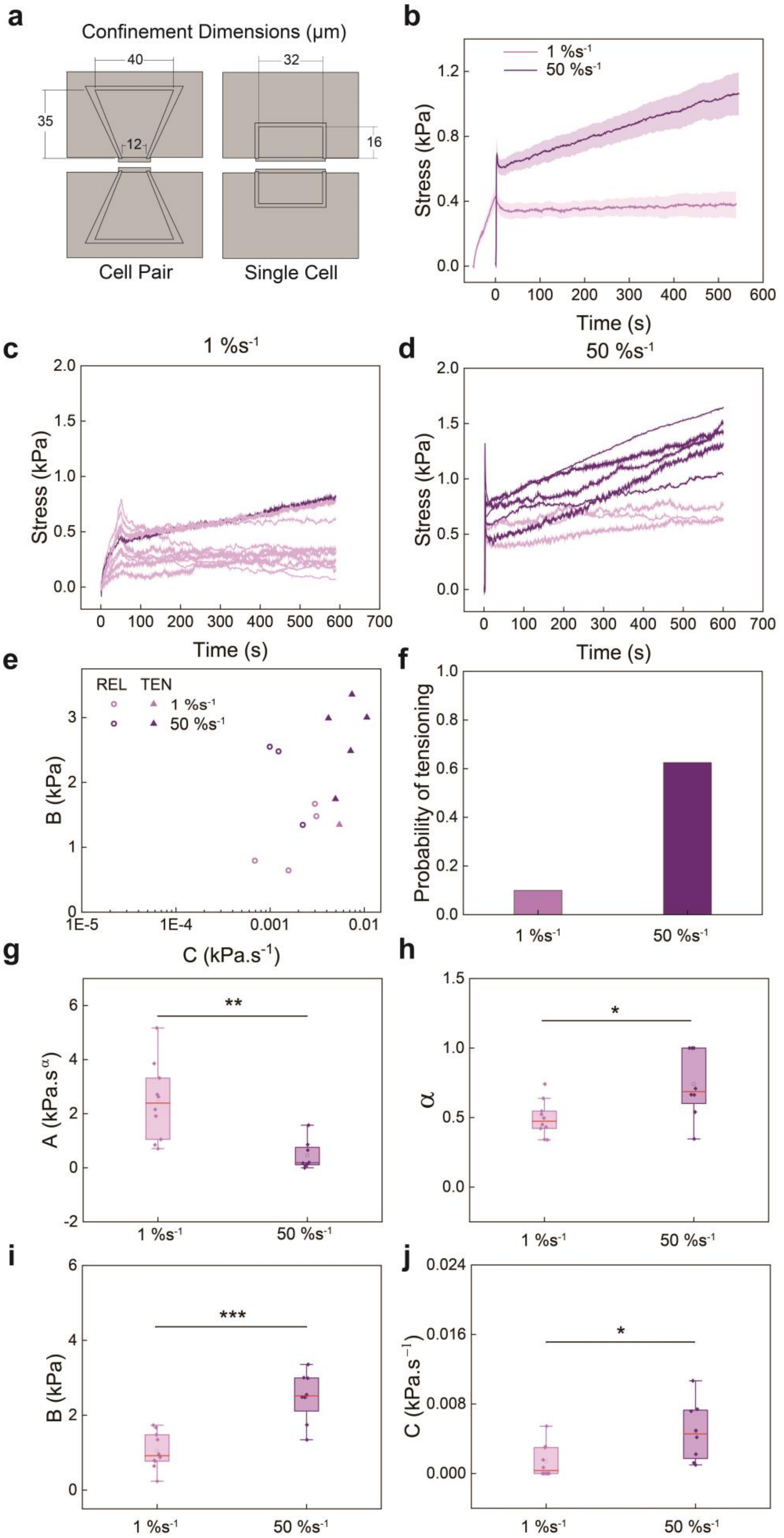
a
Confinement Dimensions (μm)
40
32
35
12
16
Cell Pair
Single Cell
b
1 %s-1
50 %s-1
Stress (kPa)
Time (s)
c
1 %s-1
d
50 %s-1
e
REL TEN
B (kPa)
C (kPa.s-1)
f
Probability of tensioning
g
A (kPa.sα)
h
α
i
j
C (kPa.s-1)
1 %s-1
50 %s-1

**Supplementary Fig. 10. The response of single cells is also strain rate dependent. a.** Illustration of the bowtie and square confinements which are used in the cell-pair and single cell studies, respectively. The area of the square confinement is equal to the area of one of the sections of the bowtie confinement. **b.** Temporal evolution of the response of single cells to 50% strain applied at 1 $\%s^{-1}$ (N = 10) and 50 $\%s^{-1}$ (N = 8) (average of all curves for each condition $\pm$ s.e.). **c-d.** Response of single cells to 50% strain applied at 1 $\%s^{-1}$ (**c**) and at 50 $\%s^{-1}$ (**d**). The curves clustered as relaxed and tensioned are shown in colors pink and purple, respectively. **e.** The B vs C values calculated from fitting individual curves for the single cell study are clustered on a semi log scale. The solid triangles represent tensioned, and the hollow circles represent the relaxed response. **f.** Probability of tensioning for single cells stretched at 1 $\%s^{-1}$ and 50 $\%s^{-1}$ strain rates. **g-j.** Box plots comparing the values of $A \cdot \varepsilon_0$ (amplitude of relaxation), $\alpha$ (power law exponent), $B \cdot \varepsilon_0$ (residual stress) and $C \cdot \varepsilon_0$ (captures the slope of the tensioning) for single cells in response to 50% strain applied at 1 $\%s^{-1}$ and 50 $\%s^{-1}$.

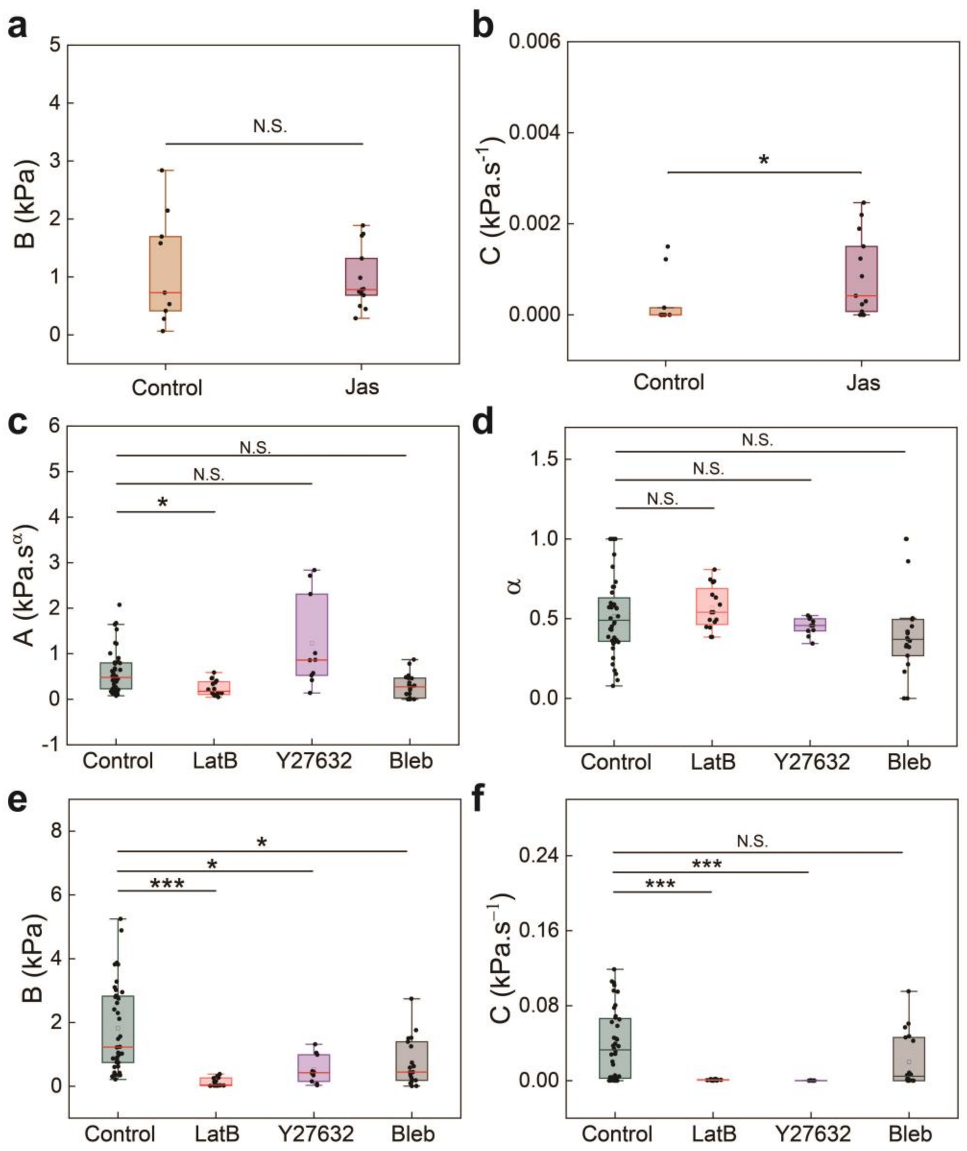


**Supplementary Fig. 11. Model parameters for drug studies.** Empirical function fitting results**. a-b.** Box plots comparing the values of the $B \cdot \varepsilon_0$ (residual stress) and $C \cdot \varepsilon_0$ (captures tensioning) for the control and Jas treated cell pairs. In this study, 125% strain was applied at 50 %s$^{-1}$. **c-f.** Box plots comparing the values of $A \cdot \varepsilon_0$ (amplitude of relaxation), $\alpha$ (power law exponent), $B$, and $C$ for the cell pairs treated with actomyosin modulators and the control group in response to 50% strain applied at 25 %s$^{-1}$.

**Table S1. Probability of tensioning.** The probability of tensioning for each condition is defined as the number of tensioned curves divided by the number of all curves.

| Experiment | | No. Relaxed curves | No. Tensioned curves | No. All curves | Probability |
|---|---|---|---|---|---|
| 25% | 0.5 $\%s^{-1}$ | 24 | 4 | 28 | 0.143 |
| | 25 $\%s^{-1}$ | 26 | 0 | 26 | 0 |
| | 50 $\%s^{-1}$ | 15 | 5 | 20 | 0.25 |
| 50% | 0.5 $\%s^{-1}$ | 19 | 0 | 19 | 0 |
| | 2.5 $\%s^{-1}$ | 12 | 0 | 12 | 0 |
| | 5 $\%s^{-1}$ | 21 | 0 | 21 | 0 |
| | 10 $\%s^{-1}$ | 20 | 10 | 30 | 0.333 |
| | 25 $\%s^{-1}$ | 14 | 26 | 40 | 0.65 |
| | 50 $\%s^{-1}$ | 7 | 24 | 31 | 0.774 |
| 125 % | 0.5 $\%s^{-1}$ | 10 | 0 | 10 | 0 |
| | 25 $\%s^{-1}$ | 7 | 0 | 7 | 0 |
| | 50 $\%s^{-1}$ | 8 | 2 | 10 | 0.2 |
| 100%, 25 $\%s^{-1}$ | One step | 12 | 0 | 12 | 0 |
| | Stretch-Hold-Stretch | 9 | 5 | 14 | 0.35 |
| Single cells | 1 $\%s^{-1}$ | 9 | 1 | 10 | 0.1 |
| | 50 $\%s^{-1}$ | 3 | 5 | 8 | 0.625 |
| 125%, 50 $\%s^{-1}$ | Control | 8 | 2 | 10 | 0.2 |
| | Jasplakinolide | 7 | 6 | 13 | 0.462 |
| 50%, 25 $\%s^{-1}$ | Control | 14 | 26 | 40 | 0.65 |
| | Blebbistatin | 12 | 4 | 16 | 0.25 |
| | Y27632 | 7 | 0 | 7 | 0 |
| | Latrunculin B | 16 | 0 | 16 | 0 |

**Supplementary Movie 1**

Comparison of cell pairs stretched at different rates, with t = 0 s defined as the point at which the maximum top island displacement is reached.

**Supplementary Movie 2**

Stress-time curve and brightfield video of two cell pairs stretched at 10 $\%s^{-1}$ to 50% strain, with one tensioning and the other relaxing as the strain is held constant.

**Supplementary Movie 3**

Time-series images of live and fixed cells stretched to 50% strain at 0.5 $\%s^{-1}$ and 50 $\%s^{-1}$ stained for F-actin.

**Supplementary Movie 4**

Time-series images of cells stretched to 50% strain at 0.5 $\%s^{-1}$ and 50 $\%s^{-1}$ and imaged for E-cadherin (green) and F-actin (red).
